# A vacuolar hexose transport is required for xylem development in the inflorescence stem of Arabidopsis

**DOI:** 10.1101/2020.12.09.417345

**Authors:** Emilie Aubry, Beate Hoffmann, Françoise Vilaine, Françoise Gilard, Patrick A.W. Klemens, Florence Guérard, Bertrand Gakière, H. Ekkehard Neuhaus, Catherine Bellini, Sylvie Dinant, Rozenn Le Hir

## Abstract

In higher plants, the development of the vascular system is controlled by a complex network of transcription factors. However, how nutrient availability in the vascular cells affects their development remains to be addressed. At the cellular level, cytosolic sugar availability is regulated mainly by sugar exchanges at the tonoplast through active and/or facilitated transport. In *Arabidopsis thaliana*, among the tonoplastic transporters, *SWEET16* and *SWEET17* genes have been previously localized in the vascular system. Here, using a reverse genetic approach, we propose that sugar exchanges at the tonoplast, regulated by SWEET16, are important for xylem cell division as revealed in particular by the decreased number of xylem cells in the *swt16* mutant and the expression of SWEET16 at the procambium-xylem boundary. In addition, we demonstrate that transport of hexoses mediated by SWEET16 and/or SWEET17 is required to sustain the formation of the xylem secondary cell wall. This result is in line with a defect in the xylem cell wall composition as measured by FTIR in the *swt16swt17* double mutant and by upregulation of several genes involved in secondary cell wall synthesis. Our work therefore supports a model in which xylem development is partially dependent on the exchange of hexoses at the tonoplast of xylem-forming cells.

## INTRODUCTION

The plant vasculature, composed of phloem, procambium/cambium, and xylem, is an elaborate system responsible for the transport of most biological compounds throughout the plant (Lucas et al., 2013). At the molecular level, vasculature development is governed by a complex network of transcription factors that are under the control of several signals, including hormones, peptides, and miRNAs (Fukuda and Ohashi-Ito, 2019; Smit et al., 2019). However, within this well-organized framework, a certain plasticity is required to adjust to cellular variations in terms of the availability of nutrients (i.e. sugars and amino acids).

Sugars, which represent the main source of energy, are required for metabolic activities, and they serve as a carbon reserve, and as intermediates for the synthesis of cell wall polysaccharides. Additionally, they have morphogenetic activity and act as primary messengers in signal transduction pathways (Sakr et al., 2018). It is therefore logical that modifications of sugar metabolism, transport or signaling can lead to multiple defects in plant growth and development (Eveland and Jackson, 2012). However, despite this central role, the role of sugar availability in the development of the vascular system in general and more specifically in heterotrophic tissues such as cambium and xylem is still elusive.

In these tissues, it has been suggested that lateral transport of sugars, coming from leakages from phloem sieve tubes, provides the sugars needed for vascular cell development (Minchin and McNaughton, 1987; Sibout et al., 2008; Spicer, 2014; Furze et al., 2018). Lateral transport is especially crucial for xylem secondary cell wall formation, since sugars are intermediate compounds in the synthesis of the cell wall polysaccharides which represent 80 % of the secondary cell wall (Marriott et al., 2016; Verbančič et al., 2018). The xylem tissue thus represents a strong sink for sugars that must be imported from surrounding tissues to serve as the source of carbon and energy. This is supported by the fact that perturbations in sugar transport at the plasma membrane of vascular cells, via SWEET or SUT transporters, affect the composition of the xylem secondary cell wall both in aspen and in Arabidopsis inflorescence stems (Mahboubi et al., 2013; Le Hir et al., 2015). Furthermore, in the Arabidopsis inflorescence stem, it has been suggested that movements of sucrose and/or hexoses towards the apoplast, mediated by SWEET11 and SWEET12, occur between the vascular parenchyma cells and the developing conducting cells to drive cell wall formation in a cell-specific manner (Dinant et al., 2019). Intercellular sugar availability seems, therefore, to play an important role in xylem development. However, the question remains open as to whether modification of sugar partitioning within the vasculature cells is also of importance.

The vacuole represents the main storage compartment for numerous primary and specialized metabolites including sugars (Martinoia, 2018). In tobacco leaves, up to 98% of hexoses are found in the vacuole (Heineke et al., 1994). In Arabidopsis leaves compartmentation of sugars is different. Sucrose is mostly present in the cytosol and the plastids while glucose and fructose are mostly found in the vacuole (Weiszmann et al., 2018). Sugar exchanges between the vacuole and the cytosol are therefore required for dynamic adjustment of the quantity of sugar needed for metabolic and signaling pathways. In herbaceous and ligneous plants, few sugar transporters have been functionally characterized at the tonoplast (Hedrich et al., 2015), and localization in the cells of the vascular system has been shown only for *SUC4/SUT4*, *ESL1*, *SWEET16* and *SWEET17* (Yamada et al., 2010; Payyavula et al., 2011; Chardon et al., 2013; Klemens et al., 2013). In *Populus*, sugar export mediated by the tonoplastic sucrose symporter PtaSUT4 is required for carbon partitioning between the source leaves and the lateral sinks (e.g. xylem) (Payyavula et al., 2011). In Arabidopsis, SWEET16 and SWEET17 transporters were localized in the roots (Guo et al., 2014) and we showed that the *SWEET16* promoter is active in the xylem parenchyma cells (Klemens et al., 2013), while the *SWEET17* promoter is active in the xylem parenchyma cells and young xylem cells of the Arabidopsis inflorescence stem (Chardon et al., 2013). Moreover high levels of *SWEET17* transcripts have been measured in the inflorescence stem, compared to other organs including roots, after 7 to 8 weeks of growth (Guo et al., 2014). *SWEET16* and *SWEET17* are therefore good candidates with which to assess whether the maintenance of sugar homeostasis between the cytosol and the vacuole influences xylem development in Arabidopsis.

In the present work, through a reverse genetic approach, we demonstrate that SWEET16 and SWEET17 have specific and overlapping roles during xylem development. In particular, we suggest that tonoplastic sugar exchanges across the procambium-xylem boundary regulated by SWEET16 are important for xylem cell proliferation. By using infrared spectroscopy and gene expression analysis, we also show that both *SWEET16* and *SWEET17* are required for correct development of the secondary cell wall of xylem cells. Finally, since glucose and fructose accumulation are observed in the inflorescence stem of the double mutant, we suggest that maintenance of hexose homeostasis through the action of SWEET16 and/or SWEET17 is important at different stages of xylem development.

## RESULTS

### Radial growth of the inflorescence stem is altered in the *swt16swt17* double mutant

To explore to what extent mutations in *SWEET16* and/or *SWEET17* impact inflorescence stem development in Arabidopsis, we used the previously described *sweet17-1* (hereafter called *swt17*) mutant line (Chardon et al., 2013) and identified two new T-DNA insertion lines in the *SWEET16* gene. The mutants *sweet16-3* (SM_3.1827) and *sweet16-4* (SM_3.30075) possess an insertion in the promoter region and in the first exon of the *SWEET16* gene respectively (Supplemental Figure 1A). They were named after the previous alleles already published (Guo et al., 2014). A full-length *SWEET16* transcript was detected by RT-PCR in the *sweet16-3* allele, while no full-length transcript could be detected in the *sweet16-4* allele (Supplemental Figure 1B and 1C). The *sweet16-4* allele (hereafter called *swt16*) was therefore deemed to be a null allele. We generated the double mutant *sweet16sweet17* (hereafter called *swt16swt17*) and confirmed by RT-PCR that both genes full length were absent in this double mutant (Supplemental Figure 1C).

Analysis of the inflorescence stems of the *swt16*, *swt17* and *swt16swt17* mutants showed that the area of stem cross-sections was significantly smaller compared to that of the wild-type (Figure 1A and B). More precisely, the stem of all mutant lines contained less xylem tissue compared with the wild type, while only the stem of *swt16swt17* displayed significantly less phloem tissue (Figure 1C-D). Additionally, the proportion of xylem or phloem per stem was calculated (Figure 1E-F). While no change in the proportion of phloem was observed in the mutants compared to the wild type (Figure 1E), a significant reduction in the xylem proportion was observed in the double *swt16swt17* mutant (Figure 1F).

**Figure 1.**
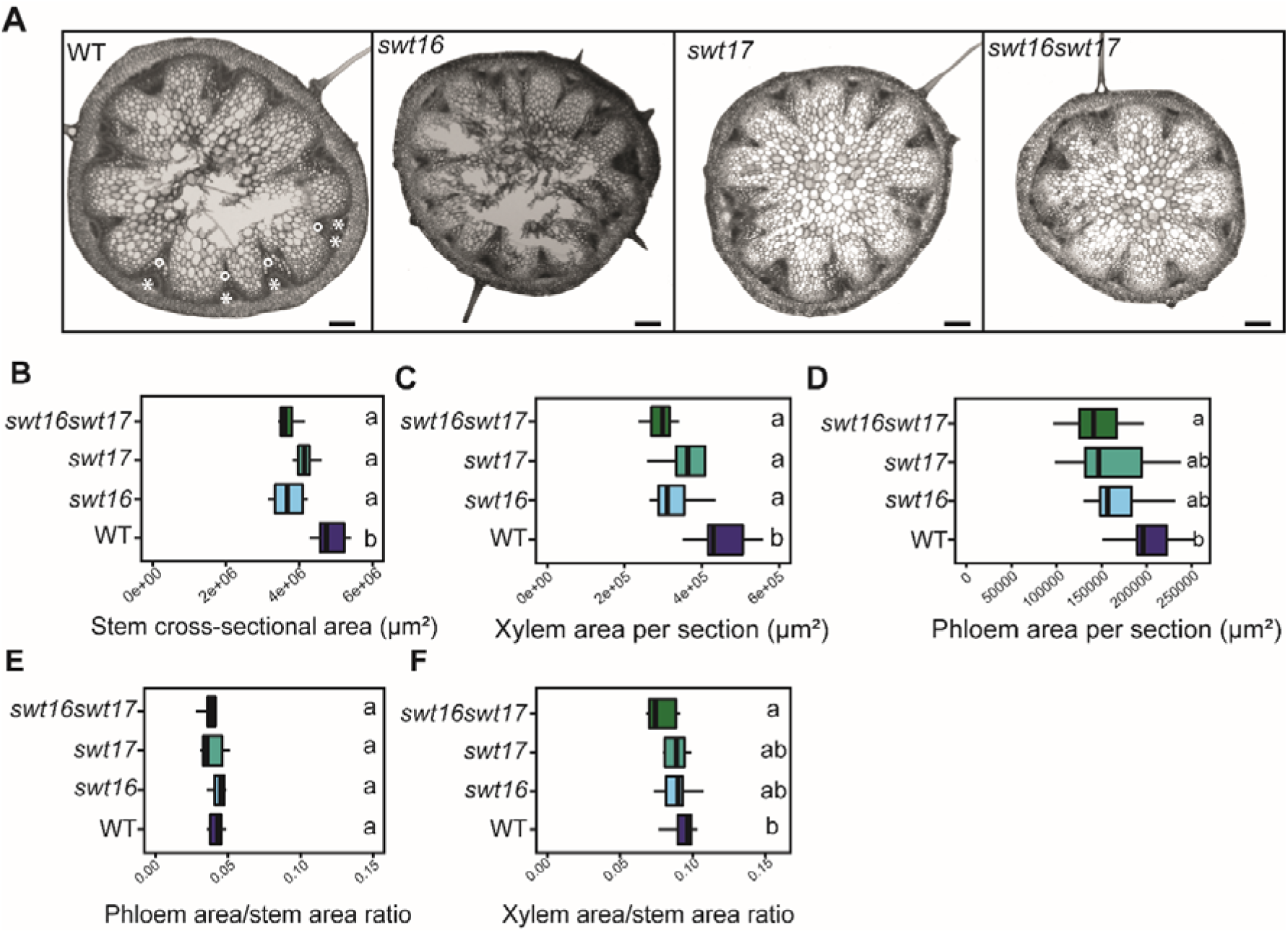
Altered development of the inflorescence stem in the *swt16swt17* double mutant. (A) Transverse sections of the basal part of the inflorescence stem of 7-week-old plants stained with FASGA solution. Stars indicate the phloem tissue while circles indicate the xylem tissue. Bars = 200 µm. (B to F) Boxplots showing the inflorescence stem cross-sectional area (B), the area occupied by xylem tissue (C) or phloem tissue (D) within a stem section, the ratio of xylem area to stem area (E) and the ratio of phloem area to stem area (F). The box and whisker plots represent values from 7 to 8 independent plants. The lines represent median values, the tops and bottoms of the boxes represent the first and third quartiles respectively, and whisker extremities represent maximum and minimum data points. A one-way analysis of variance combined with the Tukey’s comparison post-hoc test were performed. The values marked with the same letter were not significantly different from each other, whereas different letters indicate significant differences (*P* < 0.05).

To further assess the involvement of SWEET16 and/or SWEET17 in the radial growth, constructs containing N-ter GFP-fused *SWEET16* and/or *SWEET17* genomic DNA driven by their native promoter were introduced into the *swt16*, *swt17* and *swt16swt17* mutants (Supplemental Figure 2A). The GFP-SWEET17 successfully complement the phenotype of the *swt17* mutant (Supplemental Figure 2A), while GFP-SWEET16 only partially complemented the stem phenotype of the *swt16* mutant (Supplemental Figure 2A). However, full complementation of the double mutant *swt16swt17* was achieved when both translational GFP fusions were expressed (Supplemental Figure 2A).

Altogether, our results show that both SWEET16 and SWEET17 transporters are required for proper radial growth of the stem by affecting the vascular system development.

### SWEET16 and SWEET17 proteins interact physically and are expressed in the xylem during its early development

Previously, using lines expressing *pSWT:GUS* transcriptional fusions, we showed that *SWEET16* and *SWEET17* were expressed in the xylem tissue of petioles and inflorescence stems (Chardon et al., 2013; Klemens et al., 2013). To confirm the presence of SWEET16 and SWEET17 proteins in the inflorescence stem we analysed the N-ter translational GFP fusions that are complementing the mutant’s phenotype (Supplemental Figure 2A). Unfortunately, despite the phenotype complementation, we could not detect any GFP signal in these lines. Nonetheless, we obtained lines from Guo et al. (2014) expressing translational fusions between *GUS* and the C-terminus of *SWEET16* or *SWEET17* coding sequences under the control of their respective native promoter in a wild type background. We analysed the expression in three different zones: (1) a stem region where the growth was rapid, (2) a stem region where elongation growth had finished but where further thickening of the secondary cell wall was still ongoing and (3) the base of the stem, which corresponds to a mature zone (Hall and Ellis, 2013) (Figure 6).

In the region where the stem was growing rapidly SWEET16 and SWEET17 proteins are expressed in the cortex, the phloem cells, the interfascicular fibers and in the developing xylem cells (Figure 2A-B and insets). In the phloem tissue whether SWEET16 and SWEET17 were expressed in the phloem parenchyma cells and/or the companion cells will need to be further addressed by using for instance translational GFP fusions (Guo et al., 2014). In the region where secondary cell wall thickening was still ongoing and in the mature stem, SWEET16 and SWEET17 were found to be expressed in the cortex and across the phloem-procambium-xylem region (Figure 2C-F). Expression of SWEET16 and SWEET17 were also observed in young xylem cells, that could be developing xylary fibers and/or in developing xylem vessels, before extensive cell wall lignification had occurred, as shown by the weaker phloroglucinol cell wall staining (insets in Figure 2C-F). Additionally, an expression of SWEET17 was observed in xylem cells showing a lignified cell wall and situated close to the xylem vessels (Figure 2D and F). Based on their localization in the middle of the vascular bundle, this cell type are more likely to be axial parenchyma cells (Turner and Sieburth, 2002; Baghdady et al., 2006) than developing xylary fibers. Finally, SWEET17 was also expressed in xylary parenchyma cells situated at the bottom of the vascular bundle (Berthet et al., 2011) (Figure 2D and F). In addition, we also verified the expression pattern of *pSWEET16:GUS* and *pSWEET17:GUS*, used to generate the translational GFP fusions, which contains shorter promoters (1295 bp for *SWEET16* and 2004 bp for *SWEET17*) than the ones used for the translational GUS fusions (1521 bp for *SWEET16* and 2746 bp for *SWEET17*) (Supplemental Figure 2B-G). Overall we observed that the expression pattern obtained with the *pSWT:GUS* transcriptional fusions is included in that of the translational SWT:GUS fusions (Supplemental Figure 2B-G). In conclusion, SWEET16 and SWEET17 expression patterns overlap in cortex cells as well as in the phloem-procambium-xylem region and in the young xylem cells, whereas SWEET17 is specifically expressed in the xylem parenchyma cells.

**Figure 2.**
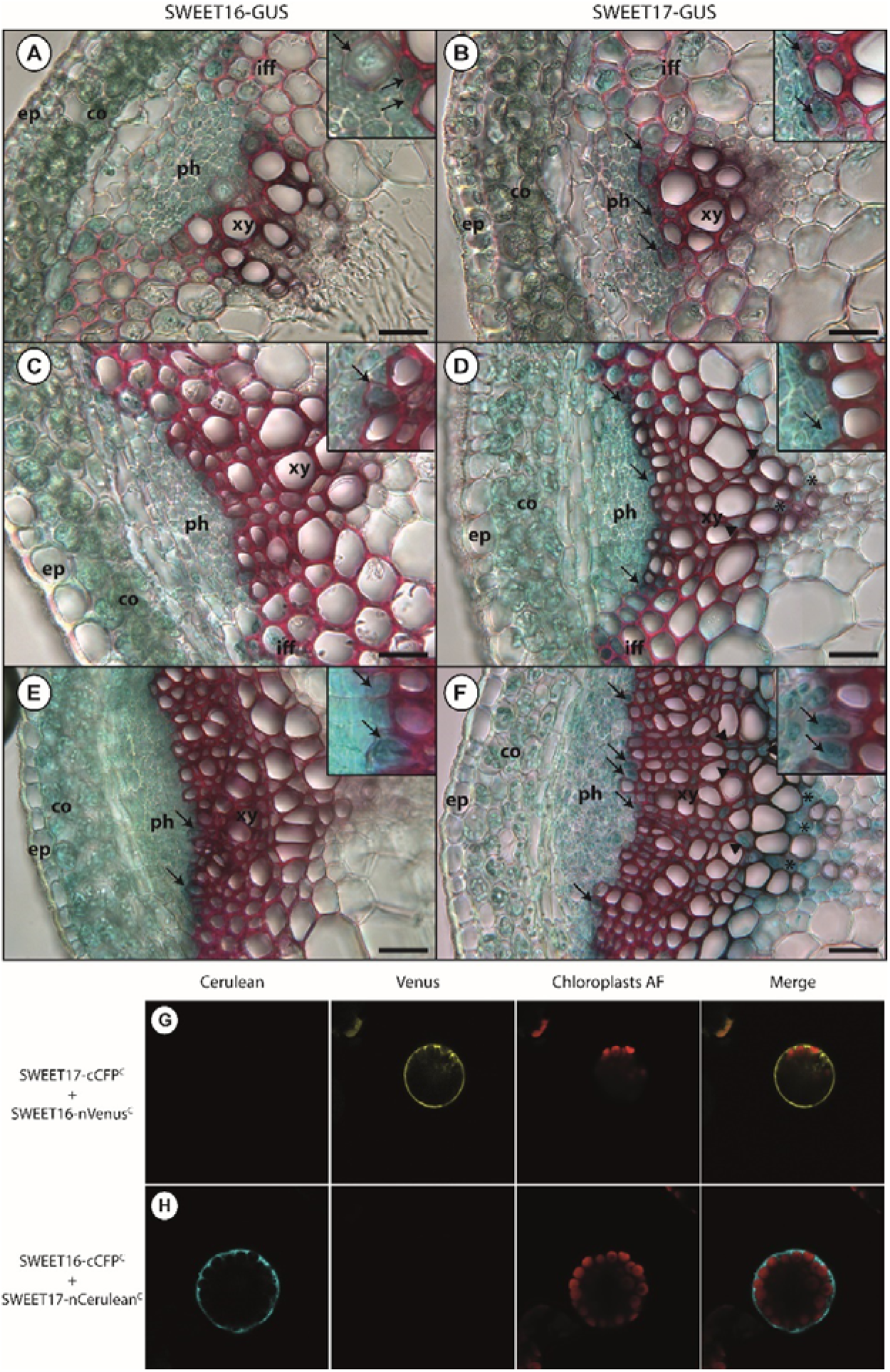
SWEET16 and SWEET17 are expressed in the xylem cells throughout the inflorescence stem development and are forming heterodimers. (A, C and E) Histochemical analysis of GUS activity in lines expression SWEET16-GUS fusion proteins driven by the *SWEET16* native promoter in sections taken at different positions in the inflorescence stem of 7-week-old plants. (B, D and F) Histochemical analysis of GUS activity in lines expression SWEET17-GUS fusion proteins driven by the *SWEET17* native promoter in sections taken at different positions in the inflorescence stem section of 7-week-old plants. Sections were taken in a stem region where the growth was still rapid (A, B and insets), in a stem region where elongation growth had finished but where thickening of the secondary cell wall was still ongoing (C, D and insets), and at the bottom of the stem, a region that corresponds to a mature stem (E, F and insets). Arrows point to cells showing blue GUS staining in developing xylem cells and arrow heads point to axial parenchyma cells and asterisks indicate xylary parenchyma cells. Lignin is colored pink after phloroglucinol staining. The intensity of the pink color is correlated with the stage of lignification of the xylary vessels. ep: epidermis; co: cortex; iff: interfascicular fibers; ph: phloem; xy: xylem. Scale bar = 50 µm. (G) Arabidopsis mesophyll protoplast expressing SWEET17-cCFP^C^ and SWEET16-nVenus^C^ interaction revealed by false color yellow, chloroplast auto-fluorescence is in false color red. (H) Arabidopsis mesophyll protoplast expressing SWEET16-cCFP^C^ and SWEET17-nCerulean^C^ interaction revealed by false color cyan, chloroplast auto-fluorescence is in false color red. Invaginations around the chloroplasts in (G) and (H) indicate that SWEET16 and SWEET17 interact at the vacuolar membrane.

It has been established that plant sugar SWEET transporters require homo or hetero-oligomerization to gain functionality (Xuan et al., 2013). Because *SWEET16* and *SWEET17* expression patterns overlap in several cells types of the inflorescence stem, we investigated whether these proteins could interact. Xuan et al. (2013) previously showed that SWEET16 and SWEET17 can form homodimers as well as heterodimers in a split ubiquitin yeast two-hybrid assay. We confirmed that SWEET16 and SWEET17 can form a heterodimer at the vacuolar membrane in a bimolecular fluorescence complementation assay in Arabidopsis mesophyll protoplasts (Figure 2G-H and Supplemental Figure 3).

### *SWEET16* but not *SWEET17* is required for proliferation of xylem cells

The localization of SWEET16 and SWEET17 in the young xylem cells prompted us to further analyzed the phenotype of the xylem tissue (Figure 3). On an independent set of plants, we first checked the robustness of the inflorescence stem phenotype and confirmed that the *swt16* and *swt17* single mutants and the *swt16swt17* double mutant consistently displayed a significantly thinner inflorescence stem compared to the wild type (Figure 3A). In addition, we confirmed our previous results that showed a significantly shorter inflorescence stem in the *swt17* mutant compared to the wild-type (Figure 3B) (Chardon et al., 2013). Interestingly, we did not observed any alteration in the inflorescence stem height in *swt16* or *swt16swt17* compared to the wild type (Figure 3B). This suggests a compensation by other sugar transporters yet to be identified in absence of SWEET16.

**Figure 3.**
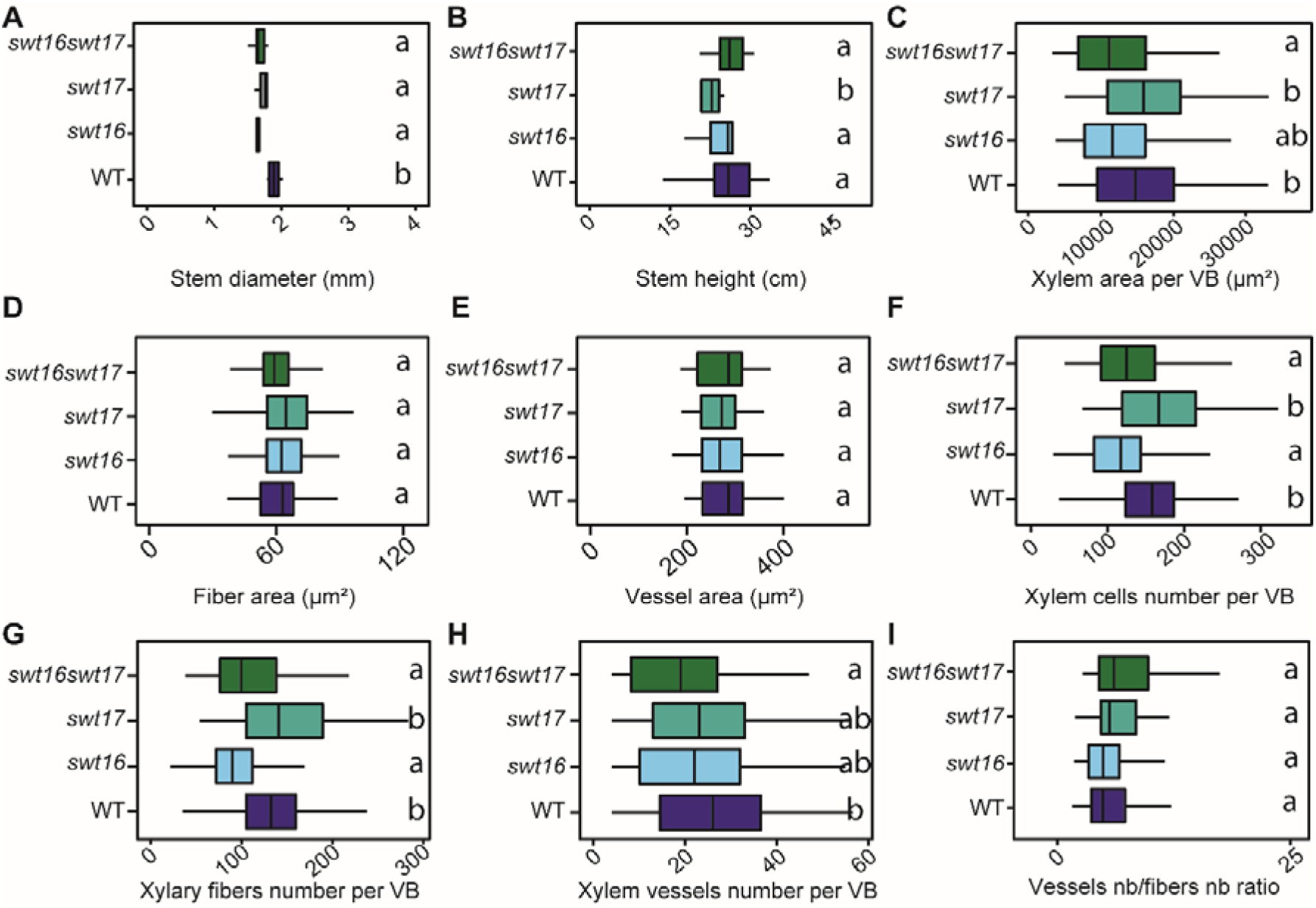
Knockout of *SWEET16* gene expression impacts the proliferation of xylem cells. (A to I) Boxplots showing the inflorescence stem height (A) and diameter (B), the cross-sectional area occupied by xylem tissue per vascular bundle (C), the average cross-sectional area of a xylary fiber (D) or of a xylem vessel (E), and the average number of xylem cells (F), of xylary fiber vessels (G), of xylem vessels (H) per vascular bundle and the ratio of vessel number to fiber number (I). The box and whisker plots represent values from 5 to 7 independent plants (A and B) or from 71, 53, 41 and 50 individual vascular bundles from wild type, *swt16*, *swt17* and *swt16swt17* respectively coming from 5-7 independent plants for each genotype (C-I). The lines represent median values, the tops and bottoms of the boxes represent the first and third quartiles respectively, and whisker extremities represent maximum and minimum data points. A one-way analysis of variance combined with the Tukey’s comparison post-hoc test were performed. The values marked with the same letter were not significantly different from each other, whereas different letters indicate significant differences (*P* < 0.05).

The xylem phenotype was then studied in more detail by counting the number of xylary fibers (cells with an area of between 5-150 µm²) and xylem vessels (cells with an area greater than 150 µm²) as well as measuring the individual cross-sectional areas within each vascular bundle (Figure 3C-I). In the vascular bundles of the *swt16* and *swt16swt17* mutants, but not *swt17*, the area occupied by the xylem tissue was significantly smaller than in the wild type (Figure 3C). These changes could result from modification of either cell size or cell number. While no changes in the size of the xylary fibers or the xylem vessels were observed in any of the genotypes analyzed (Figure 3D-E), the total number of xylem cells per vascular bundle was significantly reduced, by about 20%, in the single mutant *swt16* and the double mutant *swt16swt17* but not in *swt17* (Figure 3F). The numbers of xylary fibers and xylem vessels per vascular bundle were significantly reduced in the stem of the *swt16* single and *swt16swt17* double mutant but not in the *swt17* mutant (Figure 3G-H). The decreased number of xylary fibers was proportional to that of xylem vessels since the vessels-to-fibers ratio was the same in the wild type and the *swt16*, *swt17* and *swt16swt17* mutant lines (Figure 3I).

Overall, these results show that the single *swt16* mutant and the *swt16swt17* double mutant have the same phenotype (Figure 3 and Supplemental Table 1) and suggest that the expression of *SWEET16*, but not that of *SWEET17*, is required for correct division of xylem cells.

### *SWEET16* and *SWEET17* are required for normal secondary cell wall composition and development in xylem cells

To explore whether modifications in the vacuolar transport of sugars impact the formation of the xylem cell wall, we first exploited the transcriptomic dataset obtained from plants overexpressing a dexamethasone (DEX)-inducible version of the *VASCULAR RELATED NAC-DOMAIN PROTEIN 7* (*VND7*) gene, the master secondary wall-inducing transcription factor (Li et al., 2016). The VND7-VP16-GR plants allow the transcriptional and metabolic changes occurring during secondary cell wall formation to be studied. From the RNA-seq dataset, we extracted information related to the expression of the family of *SWEET* genes at different time points after induction of *VND7* expression (Supplemental Figure 4). Out of the 17 *SWEET* genes identified in Arabidopsis, 7 were differentially expressed during secondary cell wall formation (Supplementary Figure 4). Most interestingly, the vacuolar *SWEET2* and *SWEET17* were significantly upregulated 3 hours after DEX induction while *SWEET16* expression was upregulated 12 hours after DEX induction (Supplementary Figure 4). In contrast, the expression of genes encoding the plasma membrane localized SWEET transporters (e.g. *SWEET1*, *SWEET3*, *SWEET11* and *SWEET12*) was significantly downregulated during secondary cell wall formation (Supplementary Figure 4). Additional analysis of the dataset showed that *SWEET2* and *SWEET17* were co-regulated with genes related to cell wall synthesis (*CESA*, *SND2*, *SND3*, *MYB46*) as well as those encoding other sugar transporters localized at the tonoplast (*ESL1*) or at the plasma membrane (*STP1*, *STP13* and *PMT4*) (Supplementary Table 4). These results support the fact that, in Arabidopsis seedlings, sugar export from the vacuole to the cytosol is ongoing during secondary cell wall formation, most probably to provide sugars to be used as intermediates for cell wall formation. Since *VND7* is also expressed in xylem vessels in Arabidopsis inflorescence stems (Shi et al., 2021), we can postulate that similar sugar exchanges involving tonoplastic sugar transporters take place during secondary cell wall formation in this organ.

To assess whether *SWEET16* and *SWEET17* are indeed functionally involved in xylem secondary cell wall formation, we performed a targeted gene expression analysis including genes that are known to be part of the transcriptional network involved in stem cell proliferation/organization (*PXY*, *WOX4*) (Etchells et al., 2013), xylem cell identity (*ATHB8*) (Smetana et al., 2019), and secondary cell wall biosynthesis in vessels/fibers (*CESA4*, *CESA7*, *CESA8*, *KNAT7, MYB4, MYB43, MYB46, MYB52*, *MYB54*, *MYB58*, *MYB63*, *MYB83*, *MYB103*, *NST1*, *VND6*, *VND7*, *SND1*/*NST3*, *SND3, VNI2* and *XND1*) (Hussey et al., 2013) (Figure 4A-I). When looking at the overall transcriptional profile of the wild type and the *swt16*, *swt17* and *swt16swt17* mutants, two clusters can be identified (Figure 4A). The first cluster contains the wild type, the *swt16* and *swt17* single mutants, whereas the second includes only the *swt16swt17* double mutant. Only a subset of genes shows significantly increased expression among the different genotypes, namely *SND3*, *MYB103*, *MYB4*, *VNI2*, *SND1*, *MYB83*, *MYB54* and *MYB46* (Figure 4A), though a tendency, albeit not significant, is observed for *MYB43* (*P*=0.053) and *KNAT7* (*P*=0.091) (Figure 4A). Interestingly, all these genes belong to the transcriptional network involved in secondary cell wall biosynthesis in xylem vessels and/or in xylary fibers (for review Hussey et al., 2013). A Student’s *t*-test was then performed to compare each mutant line to the wild-type plants (Figure 4B-I). On average, a 2-fold increase in expression was measured for the genes *SND1*, *MYB46*, *VNI2*, *MYB83* and *MYB54* in the *swt16swt17* double mutant compared to the wild type (Figure 4B, C, D, F and G), while a similar tendency was observed for *MYB4*, *SND3* and *MYB103* expression (Figure 4E, H and I). Overall, these results show that in the *swt16swt17* double mutant neither stem cell maintenance nor xylem identity genes are affected, whereas secondary cell wall biosynthesis genes are deregulated.

**Figure 4.**
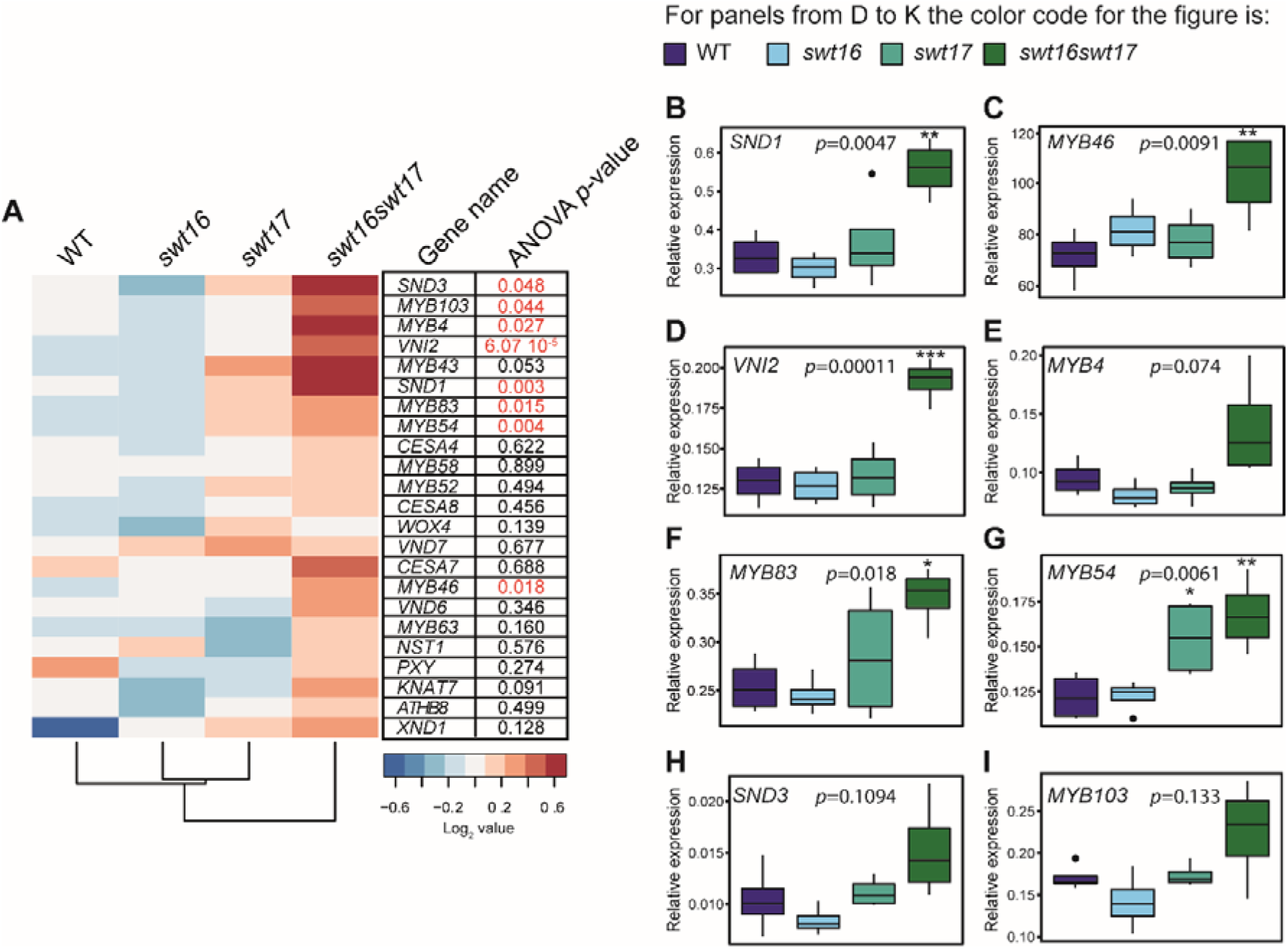
Genes involved in the development of xylem secondary cell wall are differentially regulated in the *swt16swt17* double mutant. (A to I) mRNAs were extracted from total inflorescence stems collected from 7-week-old wild-type, *swt16*, *swt17* and *swt16swt17* plants. The mRNA contents are expressed relative to those of the reference gene *UBQ5*. (A) Heatmap of expression of candidate genes involved in xylem development and secondary cell wall biosynthesis in the inflorescence stem of the wild type, *swt16*, *swt17* and *swt16swt17* mutants. The values used to build the heatmap are the mean accumulation of transcripts (*n*=4) normalized by the median value of each gene and expressed as log_2_ values. For each gene, the result of one-way ANOVA is displayed beside the heatmap. Those *p*-values below the significance threshold (*P*<0.05) are in red. (B to I) Boxplots showing the relative expression of *SND1* (B), *MYB46* (C), *VNI2* (D), *MYB4* (E), *MYB83* (F), *MYB54* (G), *SND3* (H) and *MYB103* (I). The box-and-whisker plots represent values from 4 biological replicates. The lines represent median values, the tops and bottoms of the boxes represent the first and third quartiles respectively, and the ends of the whiskers represent maximum and minimum data points. Black dots are outliers. Stars denote significant differences between the mutant line compared to the wild type according to a one-way ANOVA followed by a Dunnett post-hoc test (* *P*<0.05; ** *P*<0.01; *** *P*<0.001). The *p*-values for the comparison between wild type and *swt16swt17* are indicated on each graph. The experiment was repeated twice and gave similar results.

Next, we tested whether this transcriptional deregulation was accompanied by modifications in the cell wall composition. We used Fourier-transformed infrared spectroscopy (FTIR) on inflorescence stem cross-sections to analyze the xylem secondary cell wall composition as previously described in Le Hir et al. (2015) (Figure 5). The average spectra for all three mutants showed several differences compared to the wild-type spectra in fingerprint regions associated with cellulose, hemicelluloses and lignin (Figure 5A). The *t*-values, plotted against each wavenumber of the spectrum, showed that the mutant lines exhibited several significant negative and positive peaks (higher or lower absorbance than in the wild type) at wavenumbers associated with cellulosic and hemicellulosic polysaccharides (898 cm^-1^, 995-1120 cm^-1^, 1187 cm^-1^, 1295 cm^-1^, 1373 cm^-1^, 1401 cm^-1^, 1423 cm^-1^, 1430 cm^-1^, 1440 cm^-1^, and 1485 cm^-1^) (Åkerholm and Salmén, 2001; Kačuráková et al., 2002; Lahlali et al., 2015) (Figure 5A and B). More precisely, wavenumbers at 898 cm^-1^, associated with the amorphous region of cellulose (Kačuráková et al., 2002), and at 1430 cm^-1^, associated with crystalline cellulose (Åkerholm and Salmén, 2001), showed opposite and significant differences (Figure 5B). This suggests a potential defect in cellulose organization in the xylem secondary cell wall. Measurements of the cellulose C-O vibrations at a peak height at 1050 cm^-1^ (Lahlali et al., 2015) further indicate modifications of the cellulose composition in the cell wall of all the mutant lines (Figure 5C).

**Figure 5.**
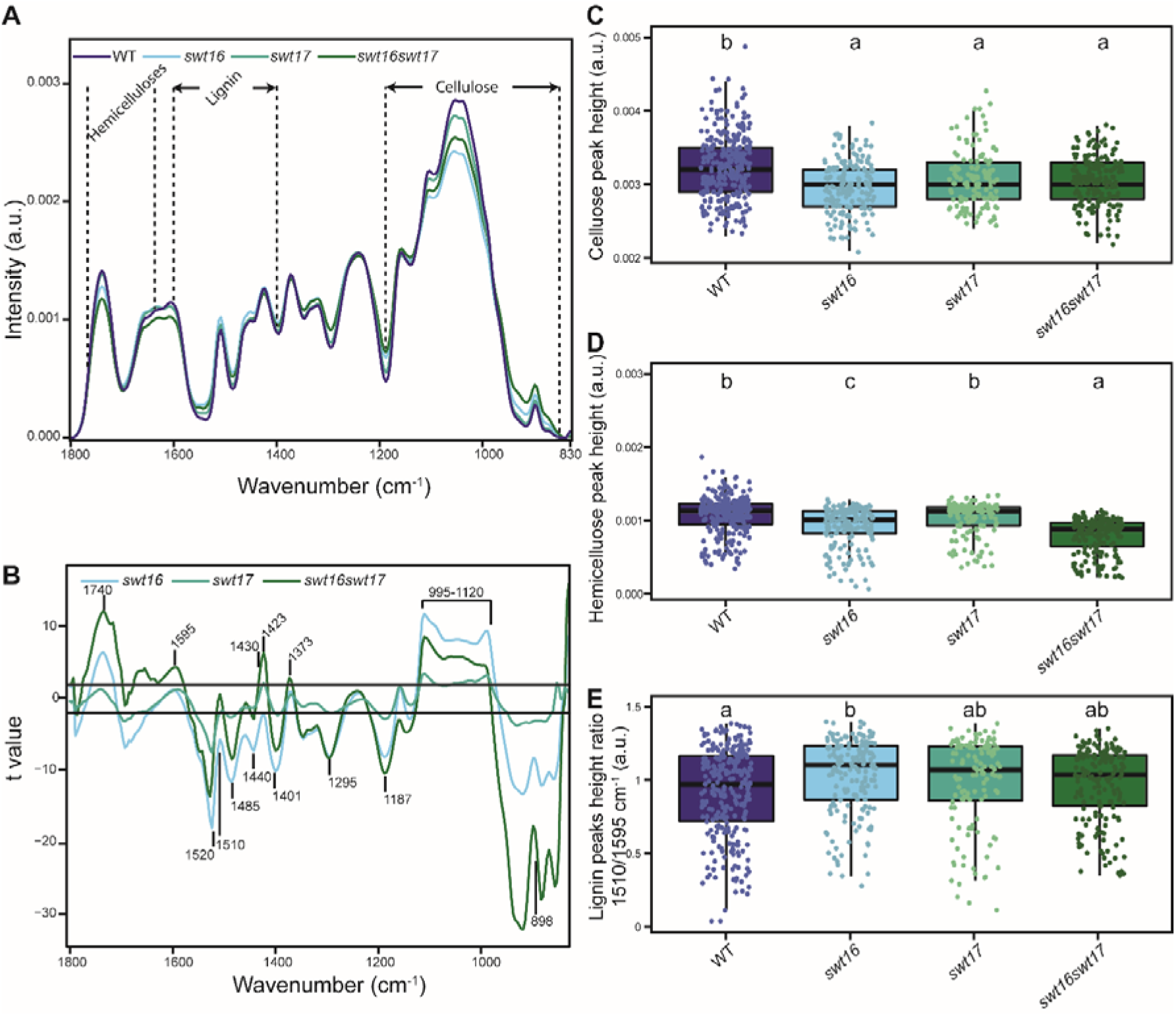
The composition of the xylem secondary cell wall is altered in the *swt16swt17* double mutant. FTIR spectra were acquired on xylem tissue from sections of the basal part of the inflorescence stem. All spectra were baseline-corrected and area-normalized in the range 1800-800 cm^-1^. (A) Average FTIR spectra were generated from 266, 170, 123 and 170 spectra for the wild type, *swt16*, *swt17* and *swt16swt17* respectively, obtained using three independent plants for each genotype. (B) Comparison of FTIR spectra obtained from xylem cells of the *swt16*, *swt17*, and *swt16swt17* mutants. A Student’s *t*-test was performed to compare the absorbances for the wild type, single and double mutants. The results were plotted against the corresponding wavenumbers. T-values (vertical axis) between −2 and +2 correspond to non-significant differences (*p*-value < 0.05) between the genotypes tested (n=3). T-values above +2 or below −2 correspond to, respectively, significantly weaker or stronger absorbances in the mutant spectra relative to the wild type. (C to E) Boxplots of the cellulose (C-O vibration band at 1050 cm^-1^) (C), hemicellulose (C-O and C-C bond stretching at 1740 cm^-1^) (D) peak height and lignin peak height ratio (1510/1595 cm^-1^) (E). The lines represent median values, the tops and bottoms of the boxes represent the first and third quartiles respectively, and the ends of the whiskers represent maximum and minimum data points. The boxplots represent values (shown as colored dots) from 266, 170, 123 and 170 spectra from the wild type, *swt16*, *swt17* and *swt16swt17* respectively, obtained from three independent plants for each genotype. A one-way analysis of variance combined with the Tukey’s comparison post-hoc test were performed. Values marked with the same letter were not significantly different from each other, whereas different letters indicate significant differences (*P* < 0.05).

The *swt16* and *swt16swt17* mutant also displayed significant variations compared to the wild type at 1740 cm^-1^ (band specific for acetylated xylan; Gou et al. 2003) and 1369 cm^-1^ (deformation of C-H linkages in the methyl group *O*-acetyl moieties; Mohebby, 2008) suggesting modifications in xylan acetylation (Figure 5A and B). Furthermore, the hemicellulose peak height at 1740 cm^-1^ (C-O and C-C bond stretching) was significantly smaller in the *swt16* mutant suggesting less acetylated xylan (Figure 5D). Although the *swt17* single mutant was not distinguishable from the wild type, the *swt16swt17* double mutant had significantly fewer acetylated xylans than the wild type and the *swt16* single mutant (Figure 5D). The lignin-associated bands at 1510 cm^-1^ (Faix, 1991), 1520 cm^-1^ (Faix, 1991; Gou et al., 2008) and 1595 cm^-1^ also exhibited significant differences in the single and double mutants compared to the wild type plants (Figure 5B). In addition we measured the lignin height peak ratio (1510/1595 cm^-1^) that can be used as a proxy for G-type lignins, to have a more detailed analysis of FTIR spectra of lignins. G-type lignins are mainly present in the cell wall of xylem vessels while G-type and S-type lignins are present in cell wall of xylary fibers (Schuetz et al., 2012; Öhman et al., 2013). We showed that the secondary cell wall of the *swt16* single mutant contains significantly more G-type lignin than that of the wild type (Figure 5E). On the other hand, only a tendency for more G-type lignin was measured for the *swt17* and the *swt16swt17* mutants which suggest that S-type lignin, also present in fibers with G-type lignin (Gorzsás et al., 2011), are responsible for the changes observed in the FTIR profiles (Figure 5A and B).

Overall, these results suggest that sugar export between the cytosol and the vacuole regulated by SWEET16 and/or SWEET17 is required to provide the intermediates needed for the synthesis of cellulosic and hemicellulosic polysaccharides.

### The hexose content is modified in the inflorescence stem of the *swt16swt17* double mutant

Assuming that SWEET16 and SWEET17 are sugar carriers, we wondered what would be the metabolic status of the inflorescence stem in the *swt16swt17* double mutant. We therefore used GC-MS to explore the global metabolomic profiles of the wild-type and the double *swt16swt17* mutant, identifying a total of 158 metabolites. In order to identify the subset of metabolites that best discriminate between the genotypes, we performed a sPLS-DA (Figure 6A). The resulting score plot clearly allows the two genotypes to be separated by the first dimension (Figure 6A). Among the metabolites selected by the sPLS-DA analysis, a subsequent *t*-test identified nine that were significantly different between wild type and the *swt16swt17* double mutant: allothreonine, benzoic acid, citraconic acid, cysteinylglycine, fructose, fumaric acid, glucose-6-phosphate, phytol and valine (Figure 6B and Supplemental Table 3). The relative quantities of benzoic acid, citraconic acid and fumaric acid (a component of the tricarboxylic cycle) were significantly reduced in the double mutant compared to the wild type (Supplemental Figure 5A, B and F). On the other hand, significant accumulation of cysteinylglycine (an intermediate in glutathione biosynthesis; (Hasanuzzaman et al., 2017), hexoses and hexose-phosphates (e.g. glucose-6-phosphate and fructose), amino acids (e.g. allothreonine and valine) and phytol (a chlorophyll component; (Gutbrod et al., 2019) was measured in the *swt16swt17* mutant compared to the wild-type stems (Supplemental Figure 5C, D, E, G, H and I). We further quantified the soluble sugars and starch content in both genotypes by enzymatic methods. Consistent with the metabolomics results, a 6-fold significant increase of fructose in the double mutant was confirmed (Figure 6C). In addition, the glucose content was significantly increased by 4-fold in the stem of the double mutant (Figure 6C), while no variation in the sucrose and starch contents was observed (Figure 6C). Interestingly, the inflorescence stem of the *swt16swt17* double mutant accumulated mostly hexoses while no significant changes in glucose or sucrose were observed in the stem of the *swt16* and *swt17* single mutants (Supplemental Figure 6A). Although it was not significant, a tendency to accumulate glucose was observed in the single mutants (Supplemental Figure 6B). A significant increase in fructose content was measured only in the *swt17* mutant compared to the wild type (Supplemental Figure 6C).

**Figure 6.**
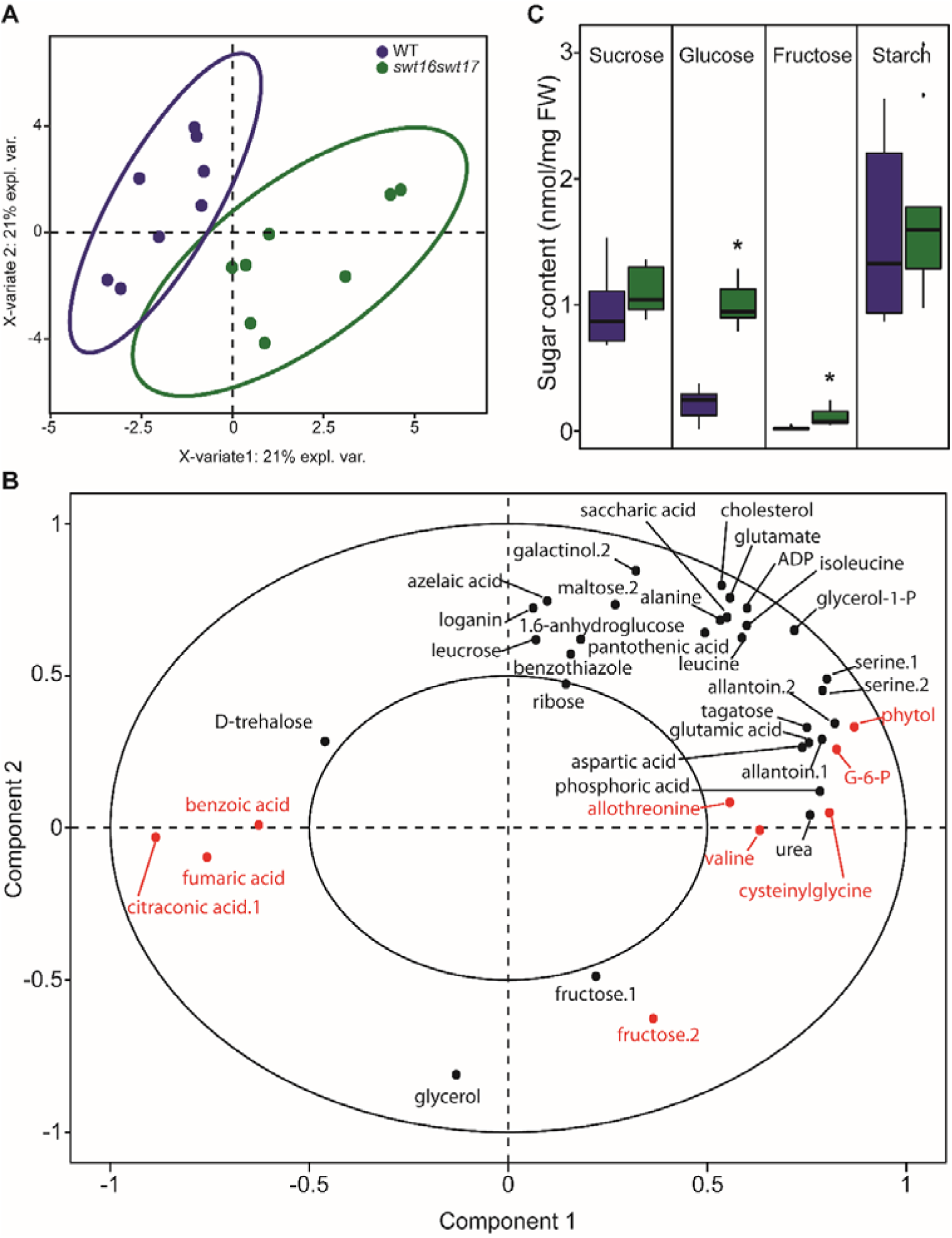
Hexoses accumulate in the inflorescence stem of the *swt16swt17* double mutant. (A and B) Multivariate analysis of the metabolomic datasets obtained from wild-type and *swt16swt17* inflorescence stems. Metabolites were extracted and analyzed by GC-MS from eight individual plants for each genotype. Plants were grown under long-day conditions for seven weeks. (A) sPLS-DA score plot for wild-type (purple) and *swt16swt17* (green) samples. The variable plot for the sPLS-DA is presented in (B) and metabolites in red are significantly different between the wild type and the *swt16swt17* mutant according to Student’s *t*-test (*P*<0.05) (Supplemental Table 3 and Supplemental Figure 3). ADP: adenosine-5-diphosphate; G-6-P: glucose-6-phosphate. (C) Boxplots showing the sucrose, glucose, fructose and starch contents of the inflorescence stems of the wild type (in purple) and the *swt16swt17* (in green) mutant grown under long-day conditions for seven weeks. The box-and-whisker plots represent values from 9 biological replicates coming from plants grown at two separate times The lines represent median values, the tops and bottoms of the boxes represent the first and third quartiles respectively, and the ends of the whiskers represent maximum and minimum data points. Black dots are outliers. Asterisks above the boxes indicate statistical differences between genotypes according to Student’s *t*-test (*P*<0.05).

## DISCUSSION

To efficiently control carbon homeostasis within a cell and to fuel the different metabolic and signaling pathways, dynamic sugar storage in the plant vacuole is critical. Over the past years, several vacuolar transporters have been identified at the molecular level (for review see Hedrich et al., 2015). Among them, SWEET16 and SWEET17 have been characterized as bidirectional tonoplastic sugar facilitators and shown to be involved in seed germination, root growth and stress tolerance (Chardon et al., 2013; Klemens et al., 2013; Guo et al., 2014; Valifard et al., 2021). In addition, the expression of both genes has been shown in the inflorescence stem’s vascular parenchyma cells, but this had not previously been explored further. In this work, we ask whether facilitated sugar transport (*via* SWEET16 and SWEET17) across the vacuolar membrane limits vascular tissue development in the inflorescence stem of Arabidopsis.

Our data show that vascular tissue development in the inflorescence stem is regulated by both SWEET16 and SWEET17 transporters. This conclusion is supported by the fact that mutations in both genes impact the inflorescence stem diameter along with the quantity of phloem and xylem tissues. These phenotypes are consistent with our analysis of lines expressing translational CDS-GUS fusions which confirmed the presence of both transporters in phloem and xylem tissues of the flower stem. Nonetheless, because SWEET16 and SWEET17 are also expressed in the root (Guo et al., 2014), the defects observed in the flower stem could also result from a disruption of sugar allocation between roots and shoot. Further experiment will be required to address the relative role of SWEET16 and SWEET17 in shoot and root parts and the consequences on development of the vascular system in the inflorescence stem.

Our data also highlight modifications of hexose homeostasis in the inflorescence stem of the different mutant lines. Although a tendency to accumulate glucose was observed in the *swt16* mutant, a significant increase in fructose was measured in the *swt17* mutant stem. Furthermore, mutations in both *SWEET16* and *SWEET17* induced somewhat specific accumulation of glucose, glucose-6-phosphate and fructose in the inflorescence stem. It has been previously shown that defects in the expression of vacuolar sugar transporters alter carbon partitioning and allocation in different organs, which is in line with our findings for the inflorescence stem (Wingenter et al., 2010; Yamada et al., 2010; Poschet et al., 2011; Chardon et al., 2013; Klemens et al., 2013; Guo et al., 2014; Klemens et al., 2014). Knowing that SWEET proteins are sugar facilitators working along the concentration gradient (Chen et al., 2010) and that at least half of the hexoses are present in the plant vacuole (Heineke et al., 1994; Weiszmann et al., 2018), we can reasonably propose that some of the hexoses are trapped inside the vacuole in the single *swt16* and *swt17* mutants and the *swt16swt17* double mutant. As a consequence, modifications in the distribution of hexose concentrations between the vacuole and the cytosol, which would impact the availability of hexoses for subsequent metabolic and signaling purposes, could be expected. Hexoses are known to favor cell division and expansion, while sucrose favors differentiation and maturation (Koch, 2004). In addition, after metabolization, hexoses and hexoses-phosphates constitute the building blocks for the synthesis of cell wall polysaccharides (Verbančič et al., 2018). Since SWEET16 and/or SWEET17 are expressed in the xylem initials, in young xylem cells and in xylem parenchyma cells, we propose that enhanced storage of vacuolar hexoses in these cells will affect different stages of xylem development.

(Pro)cambium and xylem tissues can be regarded as sinks because they rely mostly on the supply of carbohydrates from the surrounding cells to sustain their development (Sibout et al., 2008; Spicer, 2014). In aspen stem, a gradual increase in sucrose and reducing sugars, together with a rise in the activities of sugar metabolism enzymes, are observed across the cambium-xylem tissues (Roach et al., 2017). In addition, in tomato, modification of fructose phosphorylation by the fructokinase SlFRK2 leads to a defect in cambium activity (Damari-Weissler et al., 2009). Taken together, these results support the need for maintenance of sugar homeostasis in the (pro)cambium to respond to the high metabolic activity required during cell division. Our work identified SWEET16 as a player in the dividing xylem cells, acting to balance the tradeoffs between the need for sugars in the cytosol and their storage in the vacuole (Figure 7). This conclusion is supported by the fact that SWEET16 is expressed across the procambium-xylem boundary and that a mutation in *SWEET16* leads to defects in the number of xylem cells and in radial growth of the inflorescence stem. Futhermore, the expression of the gene coding for the WUSCHEL RELATED HOMEOBOX 4 (WOX4) transcription factor (Etchells et al., 2013), which is involved in cellular proliferation, was unchanged in both *swt16* and *sw16swt17* mutants. These results suggest that the defects in xylem cell division could result from reduced availability of energy and matter resources due to a reduction in sugar transport and/or from a defect in sugar signaling.

**Figure 7.**
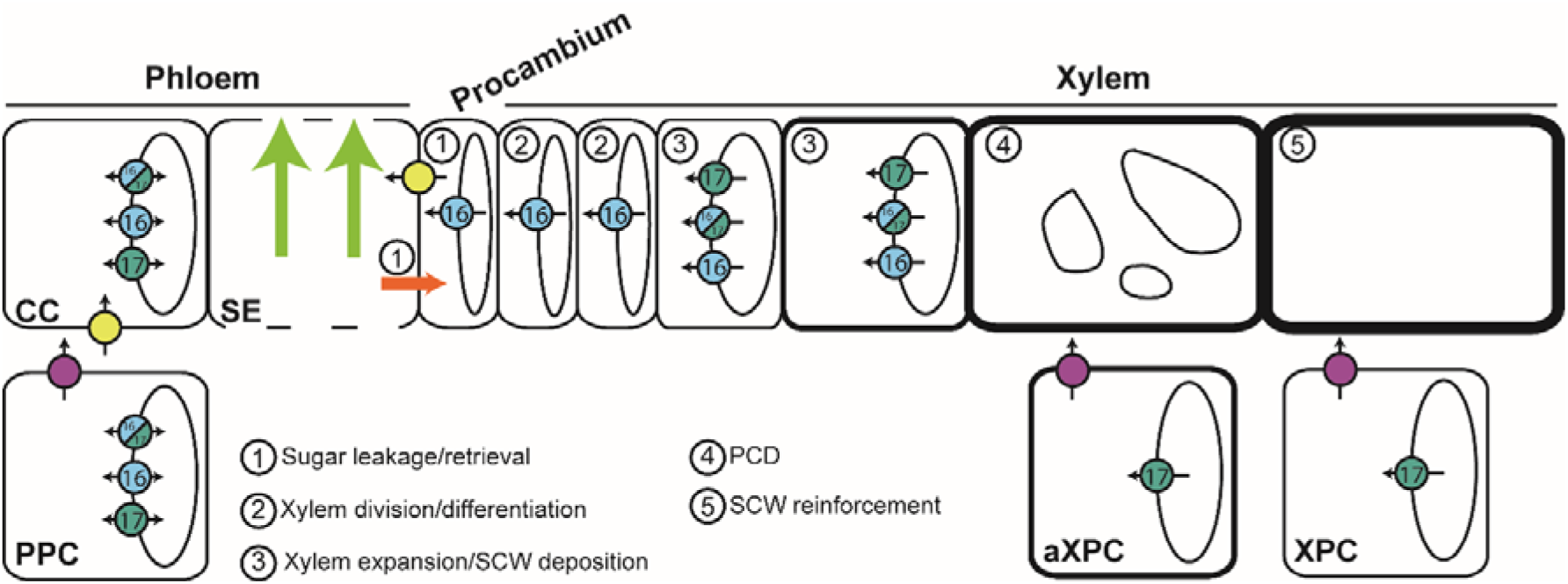
Model for the role of SWEET transporters during xylem development in Arabidopsis inflorescence stems. This model is based on the results presented in this work on SWEET16 and SWEET17 and those previously published on SWEET11, SWEET12 and SUC2 transporters (Truernit and Sauer, 1995; Chen et al., 2012; Gould et al., 2012; Le Hir et al., 2015). In the phloem tissue, sugar exchanges between cytosol and vacuole in companion cells and/or parenchyma cells are regulated by SWEET16 and SWEET17. In addition, cytosolic sucrose and hexoses present in the phloem parenchyma cells (PPC) are exported into the apoplastic space between PPC and companion cells (CC) by the sugar transporters SWEET11 and SWEET12 (fuchsia circles). Apoplastic sucrose is then imported into the CC cytosol *via* the SUC2 transporter (yellow circles) before entering the phloem sieve tubes (SE) and being transported over long distances (light green arrows). A part of these sugars leaks from the SE, most probably through plasmodesmata (orange arrow), and reaches axial sinks (e.g. procambium and xylem) while another part of the sugars is reimported inside the SE, mostly through the action of SUC2 (1). In the cells at the cambium-xylem boundary, soluble sugars are probably exported by SWEET16 (light blue respectively) into the cytosol in order to sustain the division of xylem cells (2). Given the high cytosolic sugar demand required to sustain the secondary cell wall (SCW) deposition process (3), sugars stored in the vacuole are likely exported into the cytosol through the action of SWEET16 and/or SWEET17. Interaction between SWEET16 and SWEET17 is shown as bicolor circles. After the completion of programmed cell death (PCD) and the disintegration of the vacuole (4), the SCW is still being reinforced (5) and we can assume that the sugar demand is still high. At this stage, the sugars stored in the vacuole of the xylary parenchyma cells (XPC) and the axial xylem parenchyma cells (aXPC) are likely released by SWEET17 and then exported into the apoplastic space by SWEET11 and SWEET12. Whether it is the sugars themselves or more complex cell wall sugar-derived molecules that reach the dead xylem cells remains an open question.

The reduced number of xylem vessels in the *swt16swt17* double mutant’s vascular bundle could be explained by the upregulation of *VND-INTERACTING 2* (*VNI2*), which is a repressor of the activity of the master regulator of xylem vessel differentiation VND7 (Zhong et al., 2008; Yamaguchi et al., 2010). On the other hand, the overexpression in the double mutant *swt16swt17* of SECONDARY WALL-ASSOCIATED NAC DOMAIN 1 (SND1), the master switch for fiber differentiation, would be expected to result in a shift towards increased differentiation of xylary fibers (Zhong et al., 2006), which is not consistent with the fewer fibers observed in the double mutant. Based on these results, we can assume that the increase in storage of vacuolar hexoses in the double mutant also affects xylem cell differentiation. Against this hypothesis, our results show that both xylary fibers and xylem vessels number decreased proportionately, since no change in the xylem vessels/xylary fibers ratio was measured. Consistent with this observation, the expression of the gene coding for the PHLOEM INTERCALATED WITH XYLEM (PXY) receptor, which is involved in xylem cell differentiation (Etchells et al., 2016), was not modified. Despite the upregulation of *VNI2* and *SND1* expression, which could be due to a feedback mechanism yet to be identified, these results therefore tend to suggest that no disequilibrium is occurring in xylem cell differentiation in the *swt16swt17* mutant stem. The enhanced storage of hexoses in the vacuole of the double mutant is therefore affecting the overall pattern of xylem cell division rather than xylem cell differentiation.

After cell division and differentiation, xylem cells undergo a developmental program that includes secondary cell wall formation, lignification and programmed cell death, to produce functional xylem fibers and vessels (Schuetz et al., 2012) (Figure 7). Along with the overexpression of *SND1*, the overexpression of genes encoding its downstream targets, namely MYB DOMAIN PROTEIN 46 and 83 (MYB46 and MYB83), was observed in the *swt16swt17* double mutant. Furthermore, the targets of the MYB46/MYB83 node, which positively regulates SND3 (SECONDARY WALL-ASSOCIATED NAC DOMAIN PROTEIN 3), KNAT7 (KNOTTED-LIKE HOMEOBOX OF ARABIDOPSIS THALIANA

7), MYB43 and MYB54 or/and negatively regulates MYB103, KNAT7 and MYB4, all of which are involved in the formation of the xylem secondary cell wall, are also upregulated (Hussey et al., 2013). KNAT7 directly or indirectly represses cellulose, hemicellulose and lignin biosynthetic genes (Li et al., 2012), while MYB54 is related to cellulose synthesis (Zheng et al., 2019). In Arabidopsis, MYB43 along with other MYB transcription factors regulates lignin biosynthesis (Geng et al., 2020), while its ortholog in rice is involved in the regulation of cellulose deposition (Ambavaram et al., 2011). Finally, upregulation of MYB4 in Arabidopsis results in downregulation of the lignin pathway (Jin et al., 2000), while a role for MYB103 in lignin biosynthesis has been shown in Arabidopsis stem (Öhman et al., 2013). In the *swt16wt17* double mutant, these transcriptional changes were accompanied by modifications of the secondary cell wall in terms of cellulose and hemicellulose composition. However, a single mutation in *SWEET16* or *SWEET17* was sufficient to modify the composition of the xylem cell wall without any alteration in the expression of genes involved in secondary cell wall synthesis. Our data further show that the SWEET16 and SWEET17 expression patterns overlap in xylem cells that are building a secondary cell wall and that they form a heterodimer in Arabidopsis mesophyll protoplasts. We therefore postulate that the intermediate sugars required for the synthesis of cell wall polysaccharides come in part from the vacuole unloading mediated by SWEET16 and SWEET17 homo- and heterodimers (Figure 7). Previously, it has been shown that genes encoding vacuolar sugar facilitators are up-regulated during secondary cell wall formation in xylem vessels, while sugar facilitators expressed at the plasma membrane are down-regulated (Supplementary Figure 2) (Li et al., 2016), supporting the idea that secondary cell wall formation relies on sugar export from the vacuole. In the current model of cell wall synthesis, the cytosolic catabolism of sucrose is thought to be the main source of nucleotide sugars (e.g. UDP-glucose, UDP-galactose, GDP-mannose) that act as precursors for cellulose and hemicellulose synthesis (Verbančič et al., 2018). Our data support the existence of a more complex system in which the export of vacuolar hexoses also represents a source for the synthesis of nucleotide sugars and subsequent cell wall formation (Figure 7).

Because SWEET17, a fructose-specific transporter (Chardon et al., 2013), is also expressed in the xylem parenchyma cells (including axial xylem parenchyma cells and xylary parenchyma cells), we postulate that the maintenance of fructose homeostasis within this cell type, is important and could contribute to the provision of carbon skeletons for secondary cell wall synthesis after the disappearance of the vacuole from the maturing xylem cells (Figure 7). Previously, the Arabidopsis fructokinases (FRKs), which allow the fructose phosphorylation before its metabolization by the cells, has been shown to play an important role in the vascular tissue development in hypocotyls (Stein et al., 2017). Furthermore, based on abnormal vascular cell shapes, Stein et al. (2017) proposed that FRKs may also contribute to the strength of the cell walls. This study and our results concur, therefore, to propose a link between fructose transport/metabolism and secondary wall formation that should be further explored. Within this scheme, the export of sugars in the apoplastic space between parenchyma cells and developing conducting cells could be carried out by the plasmalemmal SWEET11 and SWEET12 transporters which also expressed in the xylem parenchyma cells (Figure 7 and (Le Hir et al., 2015). Such cooperation between parenchyma cells and developing conducting cells was previously described as the “good neighbor” hypothesis in the context of H_2_O_2_ and monolignol transport (Barcelo, 2005; Smith et al., 2013; Smith et al., 2017). To further explore the importance of sugar transport between xylary parenchyma cells and developing xylem cells, more experiments, such as cell-specific complementation of the *sweet* mutant lines, will be needed in order to better comprehend the role of xylary parenchyma cells in xylem development. Additionally, it would be interesting to explore, with similar techniques, whether or not the vascular system development is impaired in the leaf petiole where both *SWEET16* and *SWEET17* genes are expressed (Chardon et al., 2013; Klemens et al., 2013). If this is the case, this would suggest a more general role of SWEET16 and SWEET17 in the xylem development.

As sink tissues, cambium and xylem rely on the lateral escape of sugars along the phloem pathway to nourish them (Sibout et al., 2008; van Bel, 2021). The localization of SWEET16 and SWEET17 at the phloem-cambium-xylem interface in the inflorescence stem along with our previous results on the localization of *SWEET11* and *SWEET12* (Le Hir et al., 2015) suggest that these SWEET transporters play a role in the radial transport (lateral transport) of sugars in order to provide energy and substrate to sustain cell division and xylem formation. However, such hypothesis needs to be tested by using for instance tissue-specific complementation to address the significance of their expression in either phloem or xylem on the global vascular system phenotype.

In conclusion, our results propose that the hexose exchanges regulated by SWEET16 are important for the division of the vascular cells while both SWEET16 and SWEET17 transporters, working as homo and/or heterodimers, are important for the secondary cell wall synthesis before the vacuole disruption. Finally, we identified SWEET17 as specifically involved in the maintenance of fructose homeostasis within the xylem parenchyma cells in order to sustain the latest phases of secondary cell wall formation. Overall, our work shows that exchange of intracellular hexoses, regulated by SWEET16 and/or SWEET17 at the tonoplast, contributes to xylem development by modulating the amounts of sugars that will be made available to the different cellular processes. However, how the cell is prioritizing the distribution of sugars among the different processes remains an open question. Although these technologies are challenging, the use of non-aqueous fractionation and metabolomics approaches (Fürtauer et al., 2019) could help in resolving subcellular sugar metabolism in a cell-specific context in mutant lines affected in sugar metabolism, transport and signaling.

## MATERIALS AND METHODS

### Plant material and growth conditions

Seeds of T-DNA insertion lines homozygous for *SWEET17* (*sweet17-1*) and *SWEET16* (*sweet16-3* and *sweet16-4*) in Col-0 background were gifts from Dr. F. Chardon and Pr. E. Neuhaus respectively. The *sweet17-1* line was previously reported to be a knock-out by Chardon et al. (2013). The *sweet16-3* (SM_3_1827) and *sweet16-4* (SM_3_30075) lines were numbered following the *sweet16-1* and *sweet16-2* lines already published by Guo et al. (2014). To verify whether *sweet16-3* and *sweet16-4* were knock-out mutants we performed RT-PCR with specific primers to amplify the full-length *SWEET16* cDNA (Supplemental Figure 1 and Supplemental Table 4). Since only the *sweet16-4* mutant turned to be a knock-out (Supplemental Figure 1B), we crossed it with *swt17-1* to obtained the double mutant *sweet16-4sweet17-1* (hereafter referred as *swt16swt17*). Homozygous plants were genotyped using gene-specific primers in combination with a specific primer for the left border of the T-DNA insertion (Supplemental Table 4).

To synchronize germination, seeds were stratified at 4°C for 48 hours and sown in soil in a growth chamber in long day conditions (16 hours day/8 hours night and 150 µE m^-2^ s^-1^) at 22/18°C (day/night temperature) with 35% relative humidity. Plants were watered with Plant-Prod nutrient solution twice a week (Fertil, https://www.fertil.fr/). For all experiments, the main inflorescence stems (after removal of lateral inflorescence stems, flowers and siliques) were harvested from seven-week old plants.

### Inflorescence stem sample preparation

For each plant, the main inflorescence stem height was measured with a ruler before harvesting a 1 to 2 cm segment taken at the bottom part of the stem. The stem segments were embedded in 8% agarose solution and sectioned with a VT100 S vibratome (Leica, https://www.leica-microsystems.com/). Some of the cross-sections were used for FT-IR analysis and the others were stained with a FASGA staining solution prepared as described in Tolivia and Tolivia (1987) for morphometric analysis of the xylem.

### Morphometric analysis of the xylem

Previously stained inflorescence stem cross-sections were imaged under an Axio Zoom V16 microscope equipped with a Plan-Neofluar Z 2.3/0.57 FWD 10.6 objective (Zeiss, https://www.zeiss.fr/microscopie/). For each section, the diameter of the inflorescence stem was measured using the Image J software package (https://imagej.nih.gov/ij/). For the same sections all the vascular bundles were photographed individually using a confocal laser scanning microscope and morphological analysis of the xylem were performed as described in Le Hir et al. (2015). For each vascular bundle, the morphological segmentation made it possible to find the number of xylem cells (xylary fibers and xylem vessels) as well as their cross-sectional areas. Cells with a cross-sectional area of between 5 to 150 µm² were considered to be xylary fibers and cells with a cross-sectional area greater than 150 µm² were considered to be xylem vessels. The sum of all xylem cell cross-sectional areas was then calculated to give the total xylem cross-sectional area. The average xylary fiber and xylem vessel area was calculated by dividing the total xylem cross-sectional area by the number of each cell type.

### GUS staining

The lines expressing pSWEET16:SWEET16-GUS and pSWEET17:SWEET17-GUS were kindly provided by Dr. Woei-Jiun Guo (National Cheng Kung University, Tainan, Taiwan) were used to assess the SWEET16 and SWEET17 expression pattern on seven-week-old plants grown under greenhouse conditions. The histochemical GUS staining was performed according to Sorin et al. (2005). Inflorescence stem subjected to GUS staining were then embedded in 8% agarose and sectioned with a Leica VT100S vibratome (Leica, https://www.leica-microsystems.com/). Sections were counterstained for lignin by phloroglucinol staining (Pradhan Mitra and Loqué, 2014). Pictures were taken using Leitz Diaplan microscope equipped with a AxioCam MRc camera and the ZEN (blue edition) software package (Zeiss, https://www.zeiss.com/).

### FT-IR analysis of the xylem secondary cell wall

The composition of the secondary cell wall of the xylem tissue was determined by Fourier Transformed Infra-red spectroscopy using an FT-IR Nicolet^TM^ iN^TM^ (Thermo Fisher Scientific, https://www.thermofisher.com). Spectral acquisitions were done in transmission mode on a 30 µm by 30 µm acquisition area targeting the xylem tissue (xylem vessels and xylary fibers) as described in Le Hir et al. (2015). Between 10 to 15 acquisition points sweeping the xylem tissue homogeneously were performed on each vascular bundle within a stem section. Three individual inflorescence stems were analyzed for each genotype. After sorting the spectra and correcting the baseline, the spectra were area-normalized and the different genotypes were compared as described in Le Hir et al. (2015). The absorbance values (maximum height) of the major cellulose, lignin and hemicellulose bands in the fingerprint region (1800–800 cm^−1^) were collected using TQ Analyst EZ edition (Thermo Fisher Scientific, https://www.thermofisher.com).

### Metabolomic analysis

The inflorescence stems of the wild-type and the *swt16swt17* double mutant were harvested in the middle of the day (8 hours after the beginning of the light period). Metabolites were extracted from 4.5 mg of lyophilized stem powder from eight individual plants and analyzed by GC-MS as described in Cañas et al. (2020). Relative concentrations of metabolites were determined relative to the internal standard ribitol, which was added after grinding the lyophilized material. Differential accumulation of metabolites was determined by one-way analysis of variance (ANOVA) and *post hoc* Tukey tests (*P<*0.05).

### Quantification of soluble sugars and starch

The main inflorescence stems of the wild type, and the *swt16*, *swt17* and *swt16swt17* mutants, were harvested in the middle of the day (8 hours after the beginning of the light period), frozen in liquid nitrogen and ground with a mortar and a pestle. Soluble sugars and starch were extracted from 50 mg of powder from an individual stem as described in Sellami et al. (2019). Depending on the experiment, 4 to 9 biological replicates were analyzed.

### RNA isolation and cDNA synthesis

RNAs were prepared from the main inflorescence stem from four 7-week-old individual plants grown as described above. Samples were frozen in liquid nitrogen before being ground with a mortar and a pestle. Powders were stored at −80°C until use. Total RNA was extracted from frozen tissue using TRIzol® reagent (Thermo Fisher Scientific, 15595-026, https://www.thermofisher.com) and treated with DNase I, RNase-free (Thermo Fisher Scientific, EN0521, https://www.thermofisher.com). cDNA was synthetized by reverse transcribing 1 µg of total RNA using RevertAid H minus reverse transcriptase (Thermo Fisher Scientific, EP0452, https://www.thermofisher.com) with 1 µl of oligo(dT)18 primer (100 pmoles) according to the manufacturer’s instructions. The reaction was stopped by incubation at 70 °C for 10 min.

### Quantitative qPCR experiment

Transcript levels were assessed for four independent biological replicates in assays with triplicate reaction mixtures by using specific primers either designed with the Primer3 software (http://bioinfo.ut.ee/primer3-0.4.0/primer3/) or taken from the literature (Supplemental Table 5). qPCR reactions were performed in a 96-well transparent plate on a Bio-Rad CFX96 Real-Time PCR machine (Bio-Rad) in 10 µl mixtures each containing 5 µl of Takyon™ ROX SYBR^®^ MasterMix dTTP Blue (Eurogentec, UF-RSMT-B0710, https://www.eurogentec.com/), 0.3 µl forward and reverse primer (30 µM each), 2.2 µl sterile water and 2.5 µl of a 1/30 dilution of cDNA. The following qPCR program was applied: initial denaturation at 95°C for 5 min, followed by thirty-nine cycles of 95°C for 10 sec, 60°C for 20 sec, 72°C for 30 sec. Melting curves were derived after each amplification by increasing the temperature in 0.5°C increments from 65°C to 95°C. The Cq values for each sample were acquired using the Bio-Rad CFX Manager 3.0 software package. The specificity of amplification was assessed for each gene, using dissociation curve analysis, by the precision of a unique dissociation peak. If one of the Cq values differed from the other two replicates by > 0.5, it was removed from the analysis. The amplification efficiencies of each primer pair were calculated from a 10-fold serial dilution series of cDNA (Supplemental Table 5). Four genes were tested as potential reference genes: *APT1* (At1g27450), *TIP41* (At4g34270), *EF1α* (At5g60390) and *UBQ5* (At3g62250). The geNorm algorithm (Vandesompele et al., 2002) was used to determine the gene most stably expressed among the different genotypes analyzed, namely *UBQ5* in this study. The relative expression level for each genotype was calculated according to the ΔCt method using the following formula: average E_t_^-Cq(of target gene in A)^/E_r_^-Cq(of reference gene in A)^, where E_t_ is the amplification efficiency of the target gene primers, E_r_ is the reference gene primer efficiency, A represents one of the genotypes analyzed.

### Production of complementation lines

For complementation of the single *sweet16-4* and *sweet17-1* and the double *sweet16-4sweet17-1* mutants, N terminal with GFP were constructed as follow. First, the coding sequence of eGFP was amplified from the pKGWFS7 plasmid (Karimi et al., 2002) with or without a stop codon and then introduced into a modified donor pENTR vector to produce pENT-GFP (w/o stop). To make the N terminal translational GFP fusions (pSWEET16:GFP-SWEET16 and pSWEET17:GFP-SWEET17), the promoters (1295 bp for *SWEET16* and 2004 bp for *SWEET17*) and genomic sequences (1863 bp for *SWEET16* and 2601 bp for *SWEET17*) were amplified separately and then cloned on each side of the GFP gene in the intermediary vector pENT-GFP by taking advantage of the restriction sites generated by PCR. All the PCR reactions were performed using Phusion High-Fidelity DNA Polymerase (Thermo Fisher Scientific, F-530S, https://www.thermofisher.com) with the primers listed in Supplemental Table 4. Donor vectors created in this way were analyzed by sequencing in order to check the reading frame of the translational fusions and the integrity of the whole genomic sequences. Destination binary vectors were then obtained by recombination, using Gateway^®^ LR Clonase II Enzyme Mix (Thermo Fisher Scientific, 11791-100, https://www.thermofisher.com), between pENTR donor vectors and pMDC99 (for pSWEET16:GFP-SWEET16) or pMDC123 (for pSWEET17:GFP-SWEET17) (Curtis and Grossniklaus, 2003). All binary vectors were introduced into *Agrobacterium tumefaciens* C58pMP90 (Koncz and Schell, 1986) by electroporation. Arabidopsis single mutants *swt16-4* and *swt17-1* as well as the double mutant *sweet16-4sweet17-1* plants were transformed by the floral dip method (Clough and Bent, 1998). Transformants were selected on hygromycin (15 mg/L) for pMDC99 constructs and/or Basta (7.5 mg/L) for pMDC123 constructs. For all constructs, three independent transgenic lines were analyzed and one representative line was selected for subsequent studies.

### BIFC assay

For the bimolecular fluorescence complementation (BiFC) assay, the full-length ORFs of *SWEET16* and *SWEET17* were amplified from cDNA with the primers given in Supplemental Table 4, either with or without their stop codons, depending on the final vector used. The ORFs were further sub-cloned into pBlueScript II SK, blunt end cut with EcoRV. The resulting vectors were checked for errors and orientation of the insert by sequencing with T3 and T7 primers. Subsequently, positive clones in the T7 orientation and the corresponding pSAT1 vectors (Lee et al., 2008) were cut with EcoRI and XhoI. *SWEET16* including the stop codon was ligated into pSAT1-cCFP-C, and *SWEET17* without the stop codon into pSAT1-nCerulean-N. Plasmid DNA of the final constructs was isolated with a PureLink™ HiPure Plasmid Filter Midiprep Kit (Invitrogen™ /Thermo Fisher Scientific) according to the manufacturer’s manual. Isolation and transfection of Arabidopsis mesophyll protoplasts were performed as described by Yoo et al. (2007). For imaging protoplasts, a Leica TCS SP5 II confocal laser scanning microscope (http://www.leica-microsystems.com) was used. All pictures were taken using a Leica HCX PL APO 63·/1.20 w motCORR CS objective with a VIS□Argon laser suitable for constructs with CFP (Cerulean, 458 nm excitation/460-490 nm collection bandwidth, laser power 90% and gain at 735 V) or YFP (Venus, 514 nm excitation/520-540 nm collection bandwidth, laser power 70% and gain at 700 V) derivates. The Chloroplast autofluorescence was imaged between 620-700 nm after excitation with the 514 nm laser line (laser power 70% and gain at 690 V).

### Statistical analysis

Differences between genotypes were assessed by a Student’s *t*-test for comparison between wild-type plants and mutant lines or by using one-way analysis of variance (ANOVA) with a Tukey HSD post-hoc test or a Dunnett post-hoc test (for analysis of the qPCR dataset). The sPLS-DA analysis was performed according to Jiang et al. (2014) and Lê Cao et al. (2011). Irrelevant variables were removed using lasso (least absolute shrinkage and selection operator) penalizations and 20 variables were selected in each dimension. The ‘mixOmics’ package (Rohart et al., 2017) was used to perform sPLS-DA. All the statistical analysis and graph production were done in RStudio (version 1.1.456) (Rstudio Team, 2015), which incorporates the R software package (version 3.5.1) (R Core Team, 2017) using ‘ggplot2’ (Wickham, 2016), ‘ggthemes’ (Arnold, 2019), ‘cowplot’ (Wilke, 2019), ‘hyperSpec’ (Beleites and Sergo, 2020) and ‘multcompView’(Graves et al., 2015).

## ACCESSION NUMBERS

Sequence data from this article can be found in the GenBank/EMBL data libraries under the following accession numbers: *SWEET16* (AT3G16690), *SWEET17* (AT4G15920). Metabolomic data can be found at https://www.ebi.ac.uk/metabolights/MTBLS2179.

## ACKNOWLEDGMENTS

The authors would like to thank Dr. Fabien Chardon and, Dr. Anne Krapp (IJPB, INRAE Versailles, France) for providing seeds of the *sweet17-1* mutant as well as Dr. Woei-Jiun Guo for kindly providing us the SWEET16 and SWEET17 translational GUS fusions. The authors also thank Dr. Grégory Mouille (IJPB, INRAE Versailles, France) for advice on FTIR dataset analysis and anonymous reviewers who helped us to improve the manuscript.

## Notes

**Funding information:** This work has benefited from the support of IJPB’s Plant Observatory technological platforms and from a French State grant (Saclay Plant Sciences, reference ANR-17-EUR-0007, EUR SPS-GSR) managed by the French National Research Agency under an Investments for the Future program (reference ANR-11-IDEX-0003-02) through PhD funding to E.A.

### Competing Interest Statement

The authors have declared no competing interest.

