## Supplemental Tables 1-5 for "A vacuolar hexose transport is required for xylem development in the inflorescence stem of Arabidopsis"

**Supplemental Tables 1 to 5 for**

### A vacuolar hexose transport is required for xylem development in the inflorescence stem of Arabidopsis.

Emilie Aubry, Beate Hoffmann, Françoise Vilaine, Françoise Gilard, Florence Guérard, Bertrand Gakière, Patrick A.W. Klemens, H. Ekkehard Neuhaus, Catherine Bellini, Sylvie Dinant and Rozenn Le Hir

Corresponding author:

**Supplemental Table 1. *p* values from pairwise comparisons (Tukey post-hoc test) between genotypes of anatomical parameters measured in the inflorescence stem xylem tissue.**

WT: wild type, *s16*: *swt16*, *s17*: *swt17* and *s16s17*: *swt16swt17*. Values in grey boxes were significantly different at the 95% confidence level.

| Compared genotypes | Stem height | Stem diameter | Total xylem area | Fiber area | Vessel area | Total xylem number | Fibers number | Vessels number | Ratio Vessels/fibers |
|---|---|---|---|---|---|---|---|---|---|
| WT - *s16* | 1 | 0.04 | 0.05 | 0.864 | 0.994 | <0.001 | <0.001 | 0.350 | 1 |
| WT - *s17* | 0.930 | 0.055 | 0.999 | 0.538 | 0.875 | 0.909 | 0.786 | 0.999 | 0.998 |
| WT - *s16s17* | 0.999 | 0.002 | 0.036 | 1 | 0.999 | 0.031 | 0.074 | 0.026 | 0.904 |
| *s16 - s17* | 0.953 | 0.980 | 0.035 | 0.998 | 0.998 | <0.001 | <0.001 | 0.680 | 0.999 |
| *s16s17 - s16* | 0.999 | 0.999 | 0.999 | 0.899 | 0.999 | 0.976 | 0.852 | 0.970 | 0.941 |
| *s16s17 - s17* | 0.899 | 0.951 | 0.025 | 0.614 | 0.990 | 0.002 | 0.002 | 0.143 | 0.999 |

**Supplemental Table 2. Genes co-regulated during the secondary cell wall formation.** Genes related to secondary cell wall formation and sugar transport were selected from differentially expressed genes associated with Cluster 21, as identified in Supplemental Table S7 in Li et al. (2016).

| AGI | Name | Biological process |
|---|---|---|
| AT1G43790 | TED6 | Cellulose biosynthesis/assembly |
| AT3G16920 | CTL2 | Cellulose biosynthesis/assembly |
| AT4G18780 | CesA8 | Cellulose biosynthesis/assembly |
| AT5G03170 | AtFLA11 | Cellulose biosynthesis/assembly |
| AT5G17420 | CesA7 | Cellulose biosynthesis/assembly |
| AT5G44030 | CesA4 | Cellulose biosynthesis/assembly |
| AT1G28470 | SND3 | Transcriptional regulation |
| AT1G62990 | IRX11 | Transcriptional regulation |
| AT1G63910 | MYB103 | Transcriptional regulation |
| AT1G66230 | MYB20 | Transcriptional regulation |
| AT2G45420 | LBD18 | Transcriptional regulation |
| AT4G00220 | LBD30 | Transcriptional regulation |
| AT4G12350 | MYB42 | Transcriptional regulation |
| AT4G22680 | MYB85 | Transcriptional regulation |
| AT4G28500 | SND2 | Transcriptional regulation |
| AT5G12870 | MYB46 | Transcriptional regulation |
| AT1G19300 | PARVUS | Xylan biosynthesis |
| AT1G27440 | IRX10/GUT2 | Xylan biosynthesis |
| AT2G28110 | FRA8 | Xylan biosynthesis |
| AT2G37090 | IRX9 | Xylan biosynthesis |
| AT3G18660 | GUX1 | Xylan biosynthesis |
| AT3G50220 | IRX15 | Xylan biosynthesis |
| AT4G33330 | GUX2 | Xylan biosynthesis |
| AT5G46340 | RWA1 | Xylan biosynthesis |
| AT5G54690 | IRX8 | Xylan biosynthesis |
| AT5G67210 | IRX15L | Xylan biosynthesis |
| AT4G15920 | SWEET17 | Fructose transport |
| AT3G14770 | SWEET2 | Glucose transport |
| AT1G08920 | ESL1 | Hexoses transport |
| AT1G11260 | STP1 | Hexoses/H+ co-transport |
| AT5G26340 | STP13 | Hexoses/H+ co-transport |
| AT2G20780 | PMT4 | Polyol/monosaccharides transport |

**Supplementary Table 3.** ***p*** **values from *t*-test of metabolites identified from the GC-MS analysis.** Values in red boxes were significantly different at the 95% confidence level. Metabolites are organized in classes according to the Golm metabolome database (http://gmd.mpimp-golm.mpg.de/Metabolites/List.aspx).

| Metabolites | *p* value | Class |
|---|---|---|
| aspartic acid 1 | 0,410 | Acid (Amino) |
| aspartic acid 2 | 0,101 | Acid (Amino) |
| Beta- alanine 1 | 0,678 | Acid (Amino) |
| DL-3-aminoisobutyric acid 2 | 0,893 | Acid (Amino) |
| DL-isoleucine 1 | 0,580 | Acid (Amino) |
| DL-isoleucine 2 | 0,345 | Acid (Amino) |
| gamma-aminobutyric acid (GABA) | 0,344 | Acid (Amino) |
| glycine | 0,550 | Acid (Amino) |
| L-alanine 2 | 0,553 | Acid (Amino) |
| L-allothreonine 2 | 0,047 | Acid (Amino) |
| L-asparagine 2 | 0,115 | Acid (Amino) |
| L-glutamic acid 2 | 0,066 | Acid (Amino) |
| L-glutamic acid 3 (dehydrated) | 0,602 | Acid (Amino) |
| L-glutamine 1 | 0,941 | Acid (Amino) |
| L-glutamine 2 | 0,224 | Acid (Amino) |
| L-glutamine 3 | 0,106 | Acid (Amino) |
| L-leucine 2 | 0,547 | Acid (Amino) |
| L-lysine 2 | 0,372 | Acid (Amino) |
| L-ornithine 2 | 0,178 | Acid (Amino) |
| L-proline 2 | 0,283 | Acid (Amino) |
| L-serine 1 | 0,096 | Acid (Amino) |
| L-serine 2 | 0,066 | Acid (Amino) |
| L-threonine 1 | 0,415 | Acid (Amino) |
| L-threonine 2 | 0,232 | Acid (Amino) |
| L-tryptophan 2 | 0,497 | Acid (Amino) |
| L-valine 2 | 0,047 | Acid (Amino) |
| norvaline 1 (pmm) | 0,973 | Acid (Amino) |
| Phenylalanine 1 | 0,162 | Acid (Amino) |
| Phenylalanine 1 | 0,124 | Acid (Amino) |
| tyrosine 2 | 0,319 | Acid (Amino) |
| citrulline 2 | 0,200 | Acid (Amino), non proteinogen |
| acetyl-L-serine 1 | 0,153 | Acid (Amino, N-acyl-) |

| Metabolites | *p* value | Class |
|---|---|---|
| 4-hydroxybenzoic acid | 0,634 | Acid (Aromatic) |
| benzoic acid (pmm) | 0,027 | Acid (Aromatic) |
| mandelic acid | 0,552 | Acid (Aromatic) |
| salicylic acid | 0,955 | Acid (Aromatic) |
| glycolic acid (sl) | 0,931 | Acid (carboxy) |
| glyoxylic acid | 0,736 | Acid (carboxy) |
| adipic acid | 0,222 | Acid (Dicarboxylic) |
| azelaic acid | 0,352 | Acid (Dicarboxylic) |
| citraconic acid 1 | 0,008 | Acid (Dicarboxylic) |
| citraconic acid 3 | 0,857 | Acid (Dicarboxylic) |
| citraconic acid 5 | 0,763 | Acid (Dicarboxylic) |
| fumaric acid | 0,039 | Acid (Dicarboxylic) |
| glutaconic acid 1 | 0,158 | Acid (dicarboxylic) |
| glutaric acid (pmm) | 0,662 | Acid (Dicarboxylic) |
| itaconic acid | 0,114 | Acid (Dicarboxylic) |
| maleic acid | 0,556 | Acid (Dicarboxylic) |
| malonic acid 1 | 0,143 | Acid (Dicarboxylic) |
| oxalic acid | 0,668 | Acid (Dicarboxylic) |
| succinic acid | 0,548 | Acid (Dicarboxylic) |
| arachidic acid | 0,779 | Acid (Fatty acid trimethylsilyl ester) |
| capric acid | 0,698 | Acid (Fatty acid trimethylsilyl ester) |
| elaidic acid | 0,787 | Acid (Fatty acid trimethylsilyl ester) |
| lauric acid | 0,505 | Acid (Fatty acid trimethylsilyl ester) |

| Metabolites | *p* value | Class |
|---|---|---|
| linoleic acid | 0,711 | Acid (Fatty acid trimethylsilyl ester) |
| myristic acid | 0,253 | Acid (Fatty acid trimethylsilyl ester) |
| Myristic Acid d27 | 0,387 | Acid (Fatty acid trimethylsilyl ester) |
| oleic acid | 0,956 | Acid (Fatty acid trimethylsilyl ester) |
| palmitic acid | 0,609 | Acid (Fatty acid trimethylsilyl ester) |
| palmitoleic acid | 0,544 | Acid (Fatty acid trimethylsilyl ester) |
| stearic acid | 0,569 | Acid (Fatty acid trimethylsilyl ester) |
| hexanoic acid (sl) | 0,277 | Acid (Fatty acid) |
| D-saccharic acid | 0,625 | Acid (Hexaric) |
| galactonic acid 2 | 0,717 | Acid (Hexonic) |
| gluconic acid 2 | 0,669 | Acid (Hexonic) |
| gluconic acid lactone 1 | 0,249 | Acid (Hexonic, lactone) |
| galacturonic acid 1 | 0,302 | Acid (Hexuronic) |
| citramalic acid | 0,460 | Acid (hydroxy ducarboxy) |
| dehydroascorbic acid 1 | 0,525 | Acid (Hydroxy) |
| dehydroascorbic acid 4 | 0,728 | Acid (Hydroxy) |
| D-malic acid | 0,795 | Acid (Hydroxy) |
| glyceric acid | 0,731 | Acid (Hydroxy) |
| L-(+) lactic acid (sl) | 0,562 | Acid (Hydroxy) |
| L-ascorbic acid | 0,162 | Acid (Hydroxy) |
| pantothenic acid 2 | 0,784 | Acid (Hydroxy) |
| quinic acid | 0,195 | Acid (Hydroxy) |
| shikimic acid | 0,744 | Acid (Hydroxy) |
| tagatose 1 | 0,051 | Acid (Hydroxy) |
| threonic acid | 0,535 | Acid (Hydroxy) |
| erythrono-1,4-lactone | 0,915 | Acid (Hydroxy, lactone) |
| alpha ketoglutaric acid | 0,114 | acid (Keto) |
| nicotinic acid | 0,308 | Acid (N-heterocycle) |
| B28pyruvic acid (sl) | 0,940 | Acid (Oxo) |
| 3,5-dimethoxy-4-hydroxycinnamique | 0,322 | Acid (Phenylpropanoic) |

| Metabolites | *p* value | Class |
|---|---|---|
| cinnamic acid | 0,847 | Acid (Phenylpropanoic) |
| ferulic acid | 0,943 | Acid (Phenylpropanoic) |
| phosphoric acid | 0,059 | Acid (Phosphate) |
| citric acid | 0,421 | Acid (Tricarboxylic) |
| isocitric acid | 0,343 | Acid (Tricarboxylic) |
| trans-aconitic acid | 0,282 | Acid (Tricarboxylic) |
| 1-decanol (decyl alcohol) | 0,114 | alcohol |
| 1-nonanol | 0,490 | alcohol |
| ethanolamine | 0,741 | Alcohol (Amino) |
| triethanolamine | 0,560 | Alcohol (Amino) |
| phytol 2 | 0,041 | Alcohol (Isoprenoid) |
| beta-glycerolphosphate | 0,540 | Alcohol (Phosphate) |
| glycerol 1-phosphate | 0,305 | Alcohol (Phosphate) |
| urea | 0,093 | Amide |
| allantoin 1 | 0,100 | Amide (N-heterocycle) |
| allantoin 3 | 0,078 | Amide (N-heterocycle) |
| putrescine | 0,891 | Amine (Poly) |
| galactinol 2 | 0,656 | Conjugate (Hexosyl, Inositol) |
| galactitol | 0,910 | Conjugate (Hexosyl, Inositol) |
| 2,3-butanediol 2 | 0,157 | misc |
| 2-hydroxypyridine | 0,942 | misc |
| 3-(methylthio)-propylamine | 0,449 | misc |
| 3-indoleacetonitrile | 0,225 | misc |
| 4-hydroxypyridine | 0,369 | misc |
| acetohydroxamic acid | 0,930 | misc |
| benzothiazole | 0,496 | misc |
| cyclohexanamine | 0,890 | misc |
| cysteinylglycine 1 | 0,040 | misc |
| homovanillic acid (HVA) | 0,839 | misc |
| isopropyl beta-D-1-thiogalacto | 0,728 | misc |
| loganin | 0,357 | misc |
| neohesperidin | 0,222 | misc |
| N-ethylglycine 2 | 0,824 | misc |
| N-methylalanine | 0,317 | misc |

| Metabolites | *p* value | Class |
|---|---|---|
| O-phosphocolamine | 0,158 | misc |
| porphine 1 | 0,736 | misc |
| tyramine | 0,691 | misc |
| adenosine 5'-diphosphate | 0,589 | Nucleoside |
| D-sorbitol | 0,539 | Polyol (Hexitol) |
| myo-inositol | 0,171 | Polyol (Inositol) |
| glycerol | 0,444 | Polyol (Triol) |
| adenine 1 | 0,357 | Purine |
| uric acid 1 | 0,711 | Purine |
| 9H-purine-6-amine | 0,133 | purine (cytokinine) |
| beta-gentiobiose 2 | 0,481 | Sugar (Disaccharide) |
| lactose 1 | 0,461 | Sugar (Disaccharide) |
| leucrose | 0,386 | Sugar (Disaccharide) |
| maltose 2 | 0,716 | Sugar (Disaccharide) |
| melibiose 1 | 0,846 | Sugar (Disaccharide) |
| Sucrose | 0,995 | Sugar (Disaccharide) |
| fructose 1 | 0,071 | Sugar (Hexose) |
| fructose 2 | 0,009 | Sugar (Hexose) |
| D (+)altrose 1 | 0,336 | Sugar (Hexose, aldose) |
| D-(+) trehalose | 0,073 | Sugar (Hexose, aldose) |
| D-glucose 1 | 0,200 | Sugar (Hexose, aldose) |
| D-glucose 2 | 0,325 | Sugar (Hexose, aldose) |
| D-mannose 1 | 0,383 | Sugar (Hexose, aldose) |
| Glucopyranose | 0,285 | Sugar (Hexose, aldose) |
| 1,5-anhydro-D-sorbitol | 0,303 | Sugar (Hexose, aldose, anhydride) |
| 1,6-anhydro-glucose | 0,435 | Sugar (Hexose, aldose, anhydride) |
| rhamnose 1 | 0,339 | Sugar (Hexose, deoxy) |
| rhamnose 2 | 0,330 | Sugar (Hexose, deoxy) |
| arabinose | 0,639 | Sugar (Pentose, aldose) |
| ribose | 0,524 | Sugar (Pentose, aldose) |

| Metabolites | *p* value | Class |
|---|---|---|
| xylose 2 | 0,591 | Sugar (Pentose, aldose) |
| xylulose | 0,773 | Sugar (Pentose, ketose) |
| D-glucose-6-phosphate 1 | 0,126 | Sugar (Phosphate) |
| D-glucose-6-phosphate 2 | 0,040 | Sugar (Phosphate) |
| fructose 6-phosphate | 0,104 | Sugar (Phosphate) |
| sedoheptulose 7-phosphate | 0,221 | Sugar (Phosphate) |
| Raffinose (pmm) | 0,931 | Sugar (Trisaccharide) |
| beta-sitosterol | 0,212 | Terpenoid (Sterols) |
| cholesterol | 0,816 | Terpenoid (Sterols) |

**Supplemental Table 4. Primers used for characterizing mutant lines.**

| Accession number | Primer name | Sequence (5’→3’) | Amplicon size (bp) | Purpose |
|---|---|---|---|---|
| At3g16690 | sweet16-3_LP | TGCAACTATGGAAATGGAAGG | 1655 | Genotyping |
| | sweet16-3_RP | GATTCAGCAAGAGCACCAAAG | | |
| | sweet16-4_LP | TGCAAATAATTTAGCAACCGC | 1742 | |
| | sweet16-4_RP | TATAAATGATCTGGGGCCATC | | |
| | SWEET16CDS+stopF | ATGGCAGACTTGAGTTTTTATGTC | 693 | Full-length PCR |
| | SWEET16CDS+stopR | TTAAGCGAGGAGAGGTTGATTT | | |
| | SWEET16CDS+stopF | ATGGCAGACTTGAGTTTTTATGTC | 693 | BIFC experiment |
| | SWEET16CDS+stopR | TTAAGCGAGGAGAGGTTGATTT | | |
| | SWEET16CDS-stopR | AGCGAGGAGAGGTTGATTT | 690 | |
| At4g15920 | sweet17-1_LP | TGATGTGAGGCCTTCCTCTT | 771 | Genotyping |
| | sweet17-1_RP | CCGTTTTGGTTGTCGTTTTT | | |
| | SWEET17CDS+stopF | ATGGCAGAGGCAAGTTTCTATATC | 726 | Full-length PCR |
| | SWEET17CDS+stopR | TTAAGAGAGGAGAGGTTCAACACG | | |
| | SWEET17CDS+stopF | ATGGCAGAGGCAAGTTTCTATATC | 726 | BIFC experiment |
| | SWEET17CDS+stopR | TTAAGAGAGGAGAGGTTCAACACG | | |
| | SWEET17CDS-stopR | AGAGAGGAGAGGTTCAACACG | 723 | |

#### Supplemental Table 5. Primers used for quantifying genes by qPCR

| Accession number | Gene name | Forward sequence (5'→3') | Reverse sequence (5'→3') | Size (bp) | Efficiency | Reference |
|---|---|---|---|---|---|---|
| At5g12870 | *MYB46* | GAATGTGAAGAAGGTGATTGGTACA | CGAAGGAACCTCAGTGTTCATCA | 150 | 73.5 | (Takeuchi et al., 2018) |
| At3g08500 | *MYB83* | GTCGCCTTCGCTGGATCAAT | AAGCCGCTTCTTCAATGTCG | 191 | 85.8 | (Shafi et al., 2019) |
| At5g61480 | *PXY* | TTCAAACCGACGAATCCATGT | TTATCCACTTGTAAAGTGTAAGCATATTCT | 85 | 95.4 | (Smetana et al., 2019) |
| At1g46480 | *WOX4* | GACAAGAACATCATCGTCACTAGACA | TTCTCCACCATTGGTTCTCTCA | 51 | 92.4 | (Smetana et al., 2019) |
| At1g32770 | *SND1/ NST3* | GCAGCAACTGGGCTAGTCTT | CCCATCGCTGCATCATAGTA | 126 | 93.6 | This study |
| At4g32880 | *AtHB8* | AACACCACTTGACCCCTCAACATCAG | CACGCAACCAACAAGGCTTATCC | 276 | 91.9 | (Carlsbecker et al., 2010) |
| At5g44030 | *CESA4* | TGCCTATGGATCGGAAAATGGA | ACGTTCTTTCCACTCCGCAT | 145 | 95.4 | (Shafi et al., 2019) |
| At5g17420 | *CESA7* | TTGTGTACGTGTCCCGTGAG | ATTTGTGAGTACGCCTGCCA | 98 | 101.1 | (Shafi et al., 2019) |
| At4g18780 | *CESA8* | AGGTCTCCCATCTGCAACAC | CTCATCGTAAGGATTGCCGC | 168 | 97.9 | (Shafi et al., 2019) |
| At1g28470 | *SND3* | TTCTTCCACCGGCCATCAAA | CTGGCGACCATAGTTGGTGT | 183 | 103.8 | (Shafi et al., 2019) |
| At1g63910 | *MYB103* | GGGAAACAGGTGGGCTCATA | TGGTAGAGGCCTCGATGGTA | 197 | 100.3 | (Shafi et al., 2019) |
| At1g62990 | *KNAT7* | GAAGCTGTTATGGCTTGCCG | TCGGTAGCAACGGACCAAAT | 190 | 95.3 | (Shafi et al., 2019) |
| At1g79180 | *MYB63* | GACAAACCCGATCTGCTGGA | CCCGAGTTCGCTTTCTAGGT | 189 | 99.8 | (Shafi et al., 2019) |
| At1g16490 | *MYB58* | AAGCGGGTTCAAAAGGTTCT | GCATCATCGTCTTTGCTTGA | 109 | 98 | This study |
| At5g13180 | *VNI2* | CTCCTTTGCCAGCTTCAATC | GGTTAGACCGGTTCCCATTT | 138 | 97.8 | This study |
| At5g64530 | *XND1* | CCCGACCTTGATCTTTACCA | CCCAATACCCATTGCTTGTC | 124 | 94.9 | This study |
| At4g39620 | *MYB4* | ACAGAGGGATTGATCCAACG | TCGACCTTTGGAGCAGAAGT | 133 | 97.9 | This study |
| At1g17950 | *MYB52* | CCGGTCGAACTGATAACGCT | ACCAATCATCCCAGTCGCAG | 131 | 103 | (Shafi et al., 2019) |
| At1g73410 | *MYB54* | AACCGAAACCCTTTCACGGA | ACGAGGCTTAGAGGTTTGGC | 174 | 87.8 | (Shafi et al., 2019) |
| At5g16600 | *MYB43* | CCATGCGCAACTTGGCAATA | CCCTTGAGCTTGTTGTGAAGC | 178 | 100.1 | (Shafi et al., 2019) |
| At3g62250 | *UBQ5* | CCAAGCCGAAGAAGATCAAG | ACTCCTTCCTCAAACGCTGA | 105 | 98.6 | This study |
