## Supplemental Figures 1-6 for "A vacuolar hexose transport is required for xylem development in the inflorescence stem of Arabidopsis"

**Supplemental Figures 1 to 6 for**

### A vacuolar hexose transport is required for xylem development in the inflorescence stem of Arabidopsis.

Emilie Aubry, Beate Hoffmann, Françoise Vilaine, Françoise Gilard, Florence Guérard, Bertrand Gakière, Patrick A.W. Klemens, H. Ekkehard Neuhaus, Catherine Bellini, Sylvie Dinant and Rozenn Le Hir

Corresponding author:

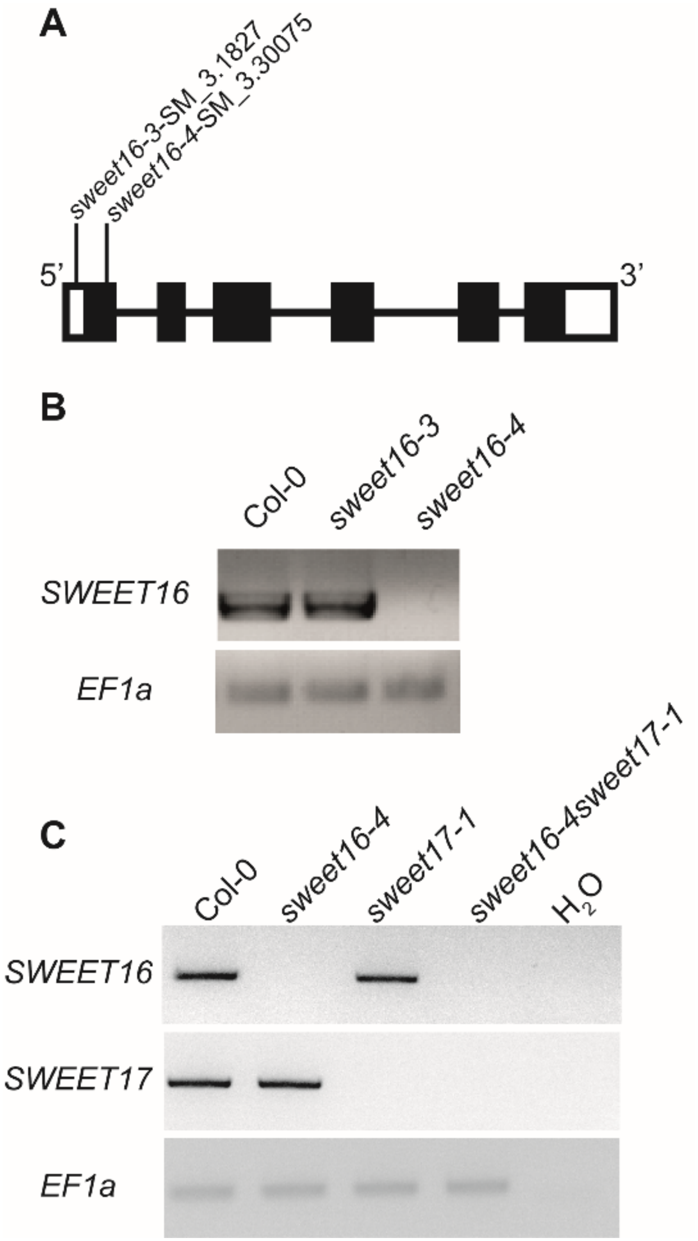


**Supplemental Figure 1. Identification of *SWEET16* insertion lines and characterization of the *sweet16-4sweet17-1* mutant line.**

(A) The position of the T-DNA insertions in *SWEET16*. White boxes represent the UTR sequences, the black boxes are the exon sequences and the black lines between the black boxes are the intron sequences.

(B) RT-PCR analysis of *SWEET16* expression in the single mutant lines *sweet16-3* and *sweet16-4*. Total RNA was isolated from 10-day-old *in vitro* seedlings and the resulting cDNA were used for amplification with primers designed between the start and stop codon of the

*SWEET16* sequence (for primers sequence see Supplementary Table 1). Expression of EF1α was used as a loading control.

(C) RT-PCR analysis of *SWEET16* and *SWEET17* expression in the floral stem of the *sweet16-4sweet17-1* mutant line. Total RNA was isolated from 7-week-old floral stem of plants grown in long-day conditions. After reverse transcription, the cDNAs were used for amplification with primers designed between the start and stop codon of the *SWEET16* and/or *SWEET17* sequences (for primers sequences see Supplemental Table 4). Expression of EF1α was used as a loading control.

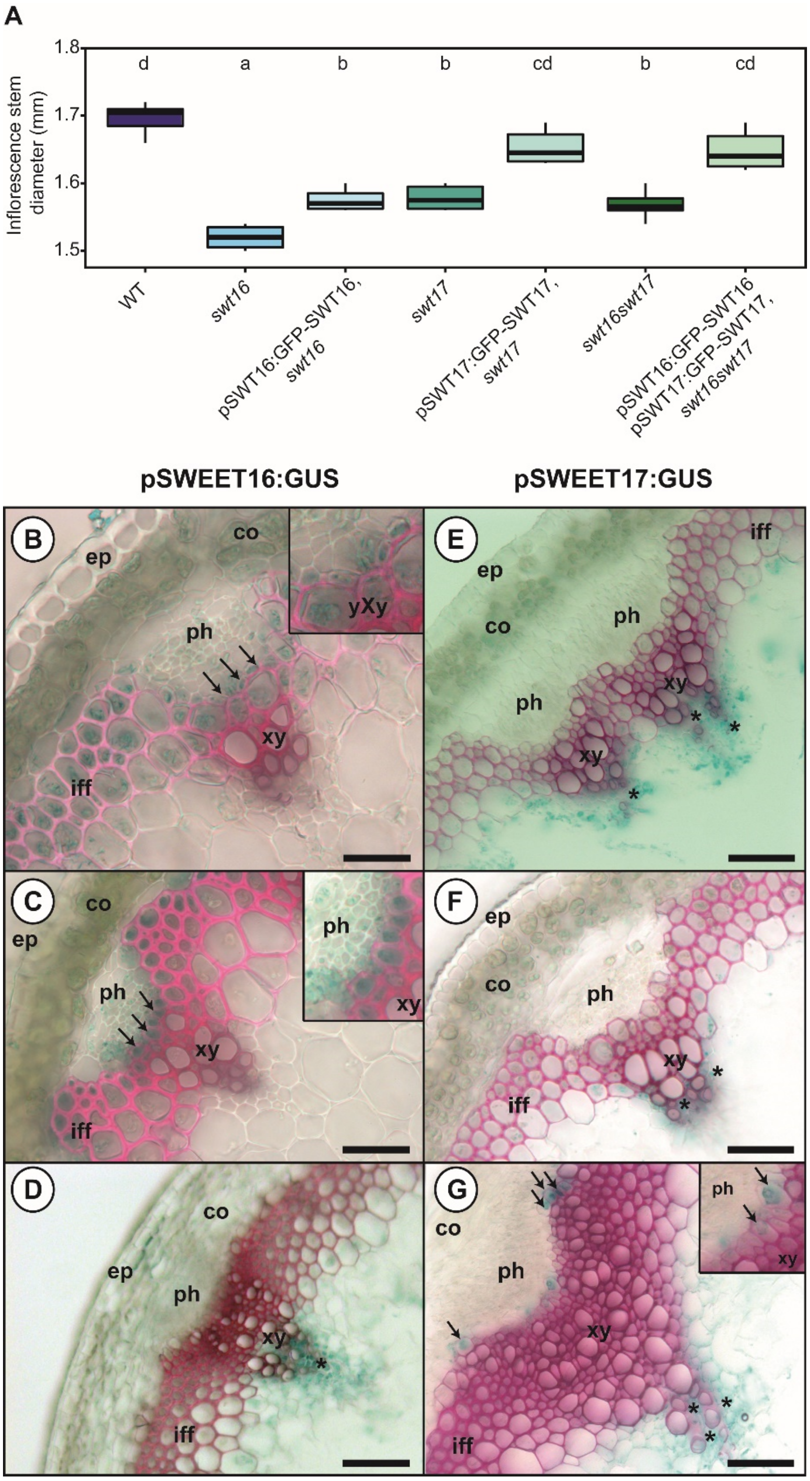


**Supplemental Figure 2. Phenotypic complementation of the *swt* mutant lines.**

(A) The diameter has been measured by a digital caliper at the bottom of the main inflorescence stem on plants grown for 7 weeks in long-day conditions. The lines represent median values, the tops and bottoms of the boxes represent the first and third quartiles, respectively, and the ends of the whiskers represent maximum and minimum data points. Values represent means from six biological replicates. A one-way analysis of variance combined with the Tukey's comparison post-test have been performed. The values marked with the same letter were not significantly different from each other, whereas different letters indicate significant differences ($P < 0.05$).

(B to D) *pSWEET16:GUS* expression pattern in sections taken at different positions in the inflorescence stem of 8-week-old plants. (E to G) *pSWEET17:GUS* expression pattern in sections taken at different positions in the inflorescence stem section of 8-week-old plants. Sections were taken in a stem region where growth was still rapid (B, E and inset), in a stem region where elongation growth had finished but where thickening of the secondary cell wall was still ongoing (C, F and inset), and at the bottom of the stem, a region that corresponds to a mature stem (D, G and inset). Arrows point to cells showing blue GUS staining and asterisks indicate xylary parenchyma cells. Lignin is colored pink after phloroglucinol staining. The intensity of the pink color is correlated with the stage of lignification of the xylary vessels. ep: epidermis; co: cortex; iff: interfascicular fibers; ph: phloem; xy: xylem. Scale bar = 50 µm.

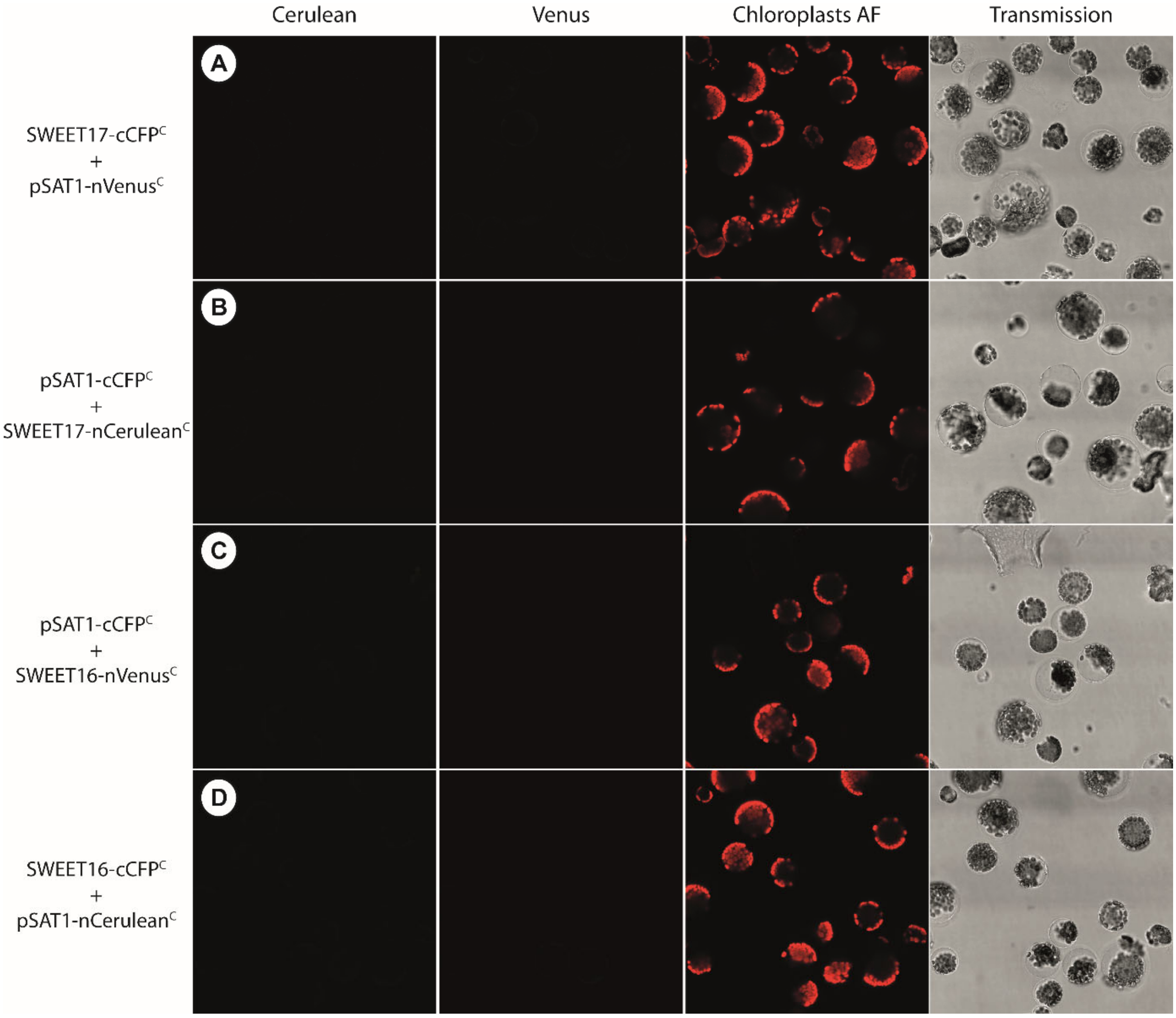


**Supplemental Figure 3. Negative controls for BIFC experiment.**

Arabidopsis mesophyll protoplasts expressing SWEET17-cCFP$^{C}$ and pSAT1-nVenus$^{C}$ (A), pSAT1-cCFP$^{C}$ and SWEET17-nCerulean$^{C}$ (B), pSAT1-cCFP$^{C}$ and SWEET16-nVenus$^{C}$ (C) and SWEET16-cCFP$^{C}$ and pSAT1-nCerulean$^{C}$ (D). No yellow (A and C) or cyan (B and D) fluorescence is reconstituted in the absence of one of the proteins. Chloroplasts autofluorescence is in false color red. The last picture of each row represents the bright field image.

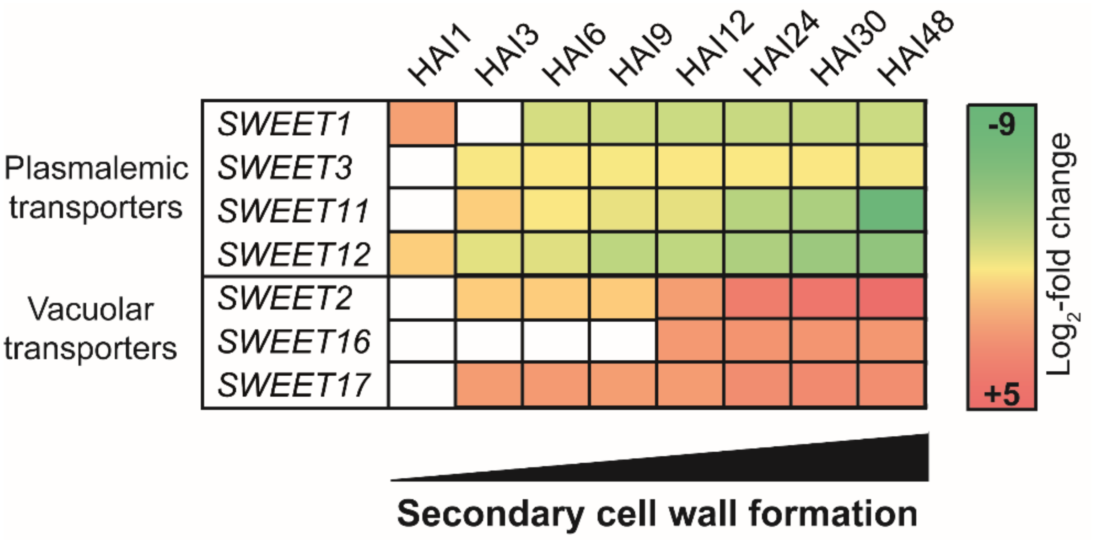


**Supplemental Figure 4. Differential expression of *SWEET* genes during the secondary cell wall formation of the xylem vessels.**

Changes in transcript levels are extracted from RNA-seq analysis performed in Li et al. (2016) and presented as $\log_2$-fold changes in comparison with the control (DMSO-treated of the DEX-inducible VND7 line) in colored boxes. HAI1-48 refer to the number of Hours After DEX Induction. White squares indicate that the gene was not differentially expressed at this time point.

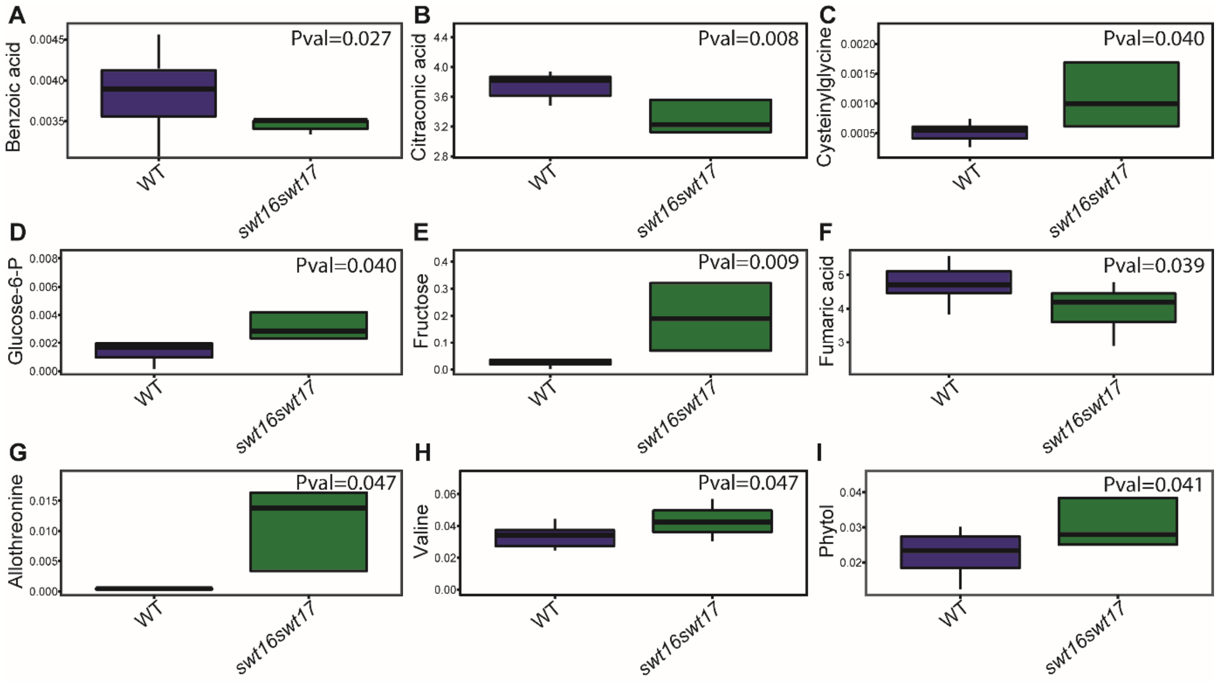


**Supplemental Figure 5. Nine out of 158 metabolites identified by GC-MS differ significantly between the wild type and the *swt16swt17* double mutant.**

Boxplots showing the relative quantity of benzoic acid (A), citraconic acid (B), cyteinylglycine (C), glucose-6-phosphate (D), fructose (E), fumaric acid (F), allothreonine (G), valine (H) and phytol (I) in the inflorescence stem of the wild type and the *swt16swt17* mutant grown in long-day conditions for seven weeks. The box-and-whisker plots represents values from 8 individual plants for each genotype. The lines represent median values, the tops and bottoms of the boxes represent the first and third quartiles, respectively, and the ends of the whiskers represent maximum and minimum data points. *P* values of the Student's *t*-test are presented directly on the graphs as well as in Supplemental Table 3.

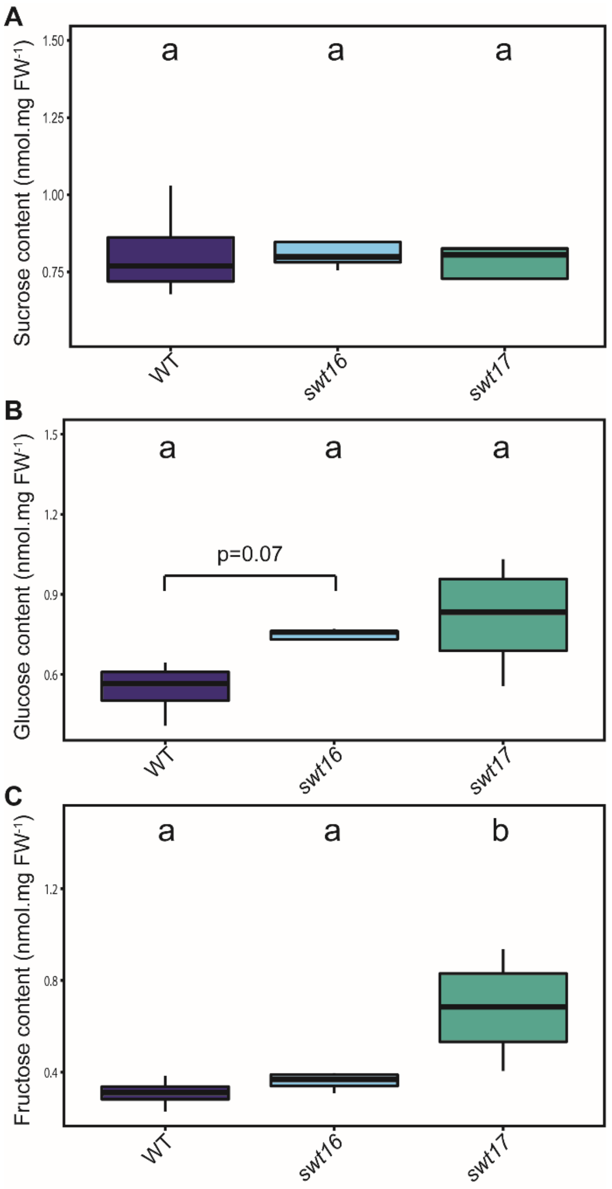


**Supplemental Figure 6. Fructose accumulation in the inflorescence stem of the *swt17* mutant.**

(A-C) Boxplots showing the sucrose (A), glucose (B) and fructose (C) contents in the inflorescence stems of the wild type, the *swt16* and the *swt17* mutants grown under long-day conditions for seven weeks. The box-and-whisker plots represents values from 4 biological replicates for each genotype. The lines represent median values, the tops and bottoms of the boxes represent the first and third quartiles, respectively, and the ends of the whiskers represent maximum and minimum data points. A one-way analysis of variance combined with the Tukey's comparison post-test have been performed. The values marked with the same letter were not significantly different from each other, whereas different letters indicate significant differences ($P < 0.05$).
